# FGF signaling induces mesoderm in members of Spiralia

**DOI:** 10.1101/2020.08.13.249714

**Authors:** Carmen Andrikou, Andreas Hejnol

## Abstract

FGF signaling is involved in mesoderm induction in deuterostomes, but not in flies and nematodes, where it has a role in mesoderm patterning and migration. However, comparable studies in other protostomic taxa are missing in order to decipher whether this mesoderm-inducing function of FGF extends beyond the lineage of deuterostomes. Here, we investigated the role of FGF signaling during mesoderm development in three species of lophophorates, a clade within the protostome group Spiralia. Our gene expression analyses show that the molecular patterning of mesoderm development is overall conserved between brachiopods and phoronids, but the spatial and temporal recruitment of transcription factors differs significantly. Moreover, inhibitor experiments demonstrate that FGF signaling is involved in mesoderm formation, morphogenetic movements of gastrulation and posterior axial elongation. Our findings suggest that the inductive role of FGF in mesoderm possibly predates the origin of deuterostomes.

## Introduction

Mesoderm is an embryonic germ layer of bilaterians that gives rise to tissues residing between the ectoderm and endoderm (Hyman, 1951; Ruppert, 1991). The way mesoderm is formed differs between embryos. For instance, deuterostomes generally form mesoderm by outpouchings of their invaginating endoderm (archenteron), a mechanism named enterocoely, which is not observed in protostomes, except for the Chaetognatha (Hertwig; Kapp, 2000; Matus et al., 2006) and Brachiopoda (Conklin, 1902; Kowalevsky, 1874; Plenk, 1913). In the remaining protostomes mesoderm originates from one or more precursor cells that are internalized during gastrulation; spiralian species can have an endodermal (e.g. the micromere 4d) and an ectodermal source (e.g. micromeres from the animal pole/anterior end of the blastopore) of mesoderm (summarized in Henry and Martindale, 1999; Kozin and Kostyuchenko, 2016; Lambert, 2008; Lyons and Henry, 2014), and ecdysozoan mesoderm originates either from internalization of vegetal endomesodermal cells (Martin-Duran and Hejnol, 2015; Sulston et al., 1983), or cells of the blastoderm (Eriksson and Tait, 2012; Hartenstein et al., 1985). Despite the differences in the embryological origin and developmental mechanisms, the molecular patterning of mesoderm induction and differentiation into various organs shares similarities between bilaterians (Amin et al., 2009; Amin et al., 2010; Andrikou et al., 2013; Chiodin et al., 2013; Fritzenwanker et al., 2014; Grifone et al., 2005; Harfe et al., 1998; Hinman and Degnan, 2002; Imai et al., 2004; Kozin et al., 2016; Kozmik et al., 2007; Mahlapuu et al., 2001; Mankoo et al., 1999; Materna et al., 2013; Nederbragt et al., 2002; Osborne et al., 2018; Passamaneck et al., 2015; Perry et al., 2015; Rudnicki et al., 1993; Sandmann et al., 2007; Schubert et al., 2003; Shimeld et al., 2010; Zaffran et al., 2001). This molecular conservation has been commonly used as an argument for the homology of this germ layer (Burton, 2008; Lartillot et al., 2002; Martindale et al., 2004; Seipel and Schmid, 2005; Technau and Scholz, 2003). Apart of shared sets of transcriptions factors, conserved signaling cascades are also involved in different steps of mesoderm development such as FGF, Notch and BMP (Good et al., 2004; Itoh and Ornitz, 2004; Sweet et al., 1999; Wijesena et al., 2017; Winnier et al., 1995). Among the aforementioned, FGF signaling is of particular interest due to its proposed ancestral role in mesoderm induction in deuterostomes (Fan et al., 2018; Green et al., 2013).

Functional studies have demonstrated that this signal is required for posterior mesoderm formation in vertebrates (Amaya et al., 1993; Draper et al., 2003; Fletcher et al., 2006; Fletcher and Harland, 2008; Yamaguchi et al., 1994), anterior mesoderm formation in cephalochordates (Bertrand et al., 2011), mesenchyme induction and formation of notochord, TVC and tail muscle in tunicates (Davidson et al., 2006; Imai et al., 2002; Kim and Nishida, 2001; Yasuo and Hudson, 2007), mesoderm induction in hemichordates (Fan et al., 2018; Green et al., 2013) and myoblast formation in sea urchins (Andrikou et al., 2015). Outside deuterostomes, however, studies addressing the role of FGF in mesoderm development are scarce. The only available data among protostome taxa concerns the two well-studied ecdysozoans *Drosophila melanogaster* and *Caenorhabditis elegans*, in which FGF is involved in mesoderm patterning and migration but not in induction (Beiman et al., 1996; Burdine et al., 1998; DeVore et al., 1995; Kadam et al., 2009; Lo et al., 2008; McMahon et al., 2010; Photos et al., 2006; Stathopoulos et al., 2004; Sun and Stathopoulos, 2018; Wilson et al., 2005). A question therefore emerges as to whether the mesoderm-inducing role of FGF originated in deuterostomes, or predated deuterostomes and got lost in the lineage of ecdysozoans (Fig. 1a). To gain insight into the ancestral role of FGF signaling for mesoderm development, data from other protostomes and, in particular, members of the Spiralia, are therefore needed.

**Fig. 1.**
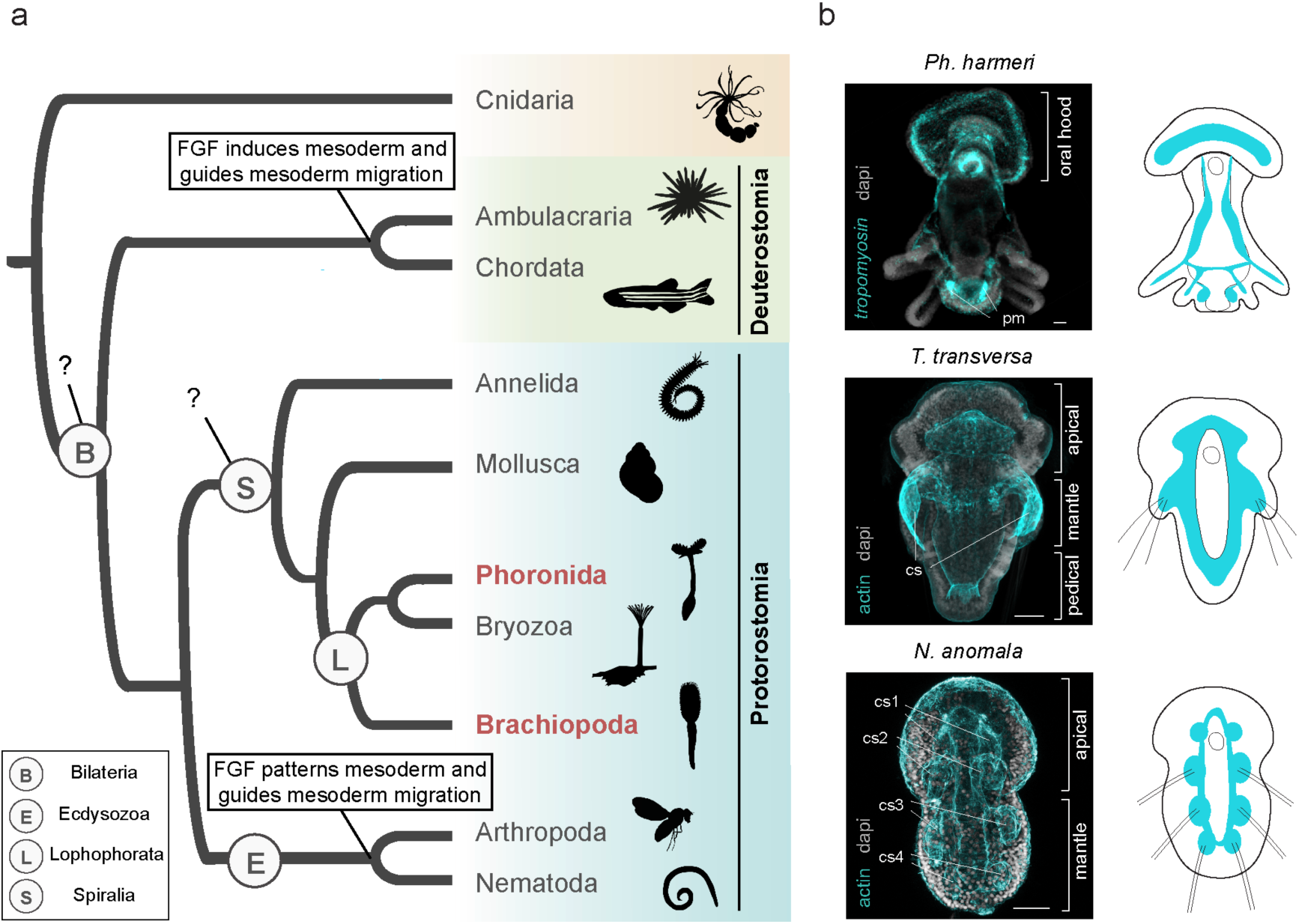
The distinct roles of FGF signaling in mesoderm development among bilaterians. (a) FGF signaling plays pivotal roles in mesoderm formation and migration in deuterostomes; however, in protostomes the information is restricted in members of ecdysozoans, where it acts in mesoderm patterning and migration. Animal illustrations are taken from phylopic.org (CC BY 3.0). (b) Mesoderm morphology in larvae of three representative lophophorate species; the brachiopods *Terebratalia transversa* and *Novocrania anomala*, and the phoronid *Phoronopsis harmeri.* Next to the images, schematic representations are shown. Drawings are not up to scale. In brachiopods mesoderm is stained by immunohistochemistry against actin and in *Ph. harmeri* mesoderm is stained by *tropomyosin* gene expression. Every fluorescent image is a full projection of merged confocal stacks and nuclei are stained with DAPI. Anterior to the top. cs, coelomic sac; pm, posterior mesoderm. Scale bar: 20 um.

Lophophorates belong to the lineage of Spiralia, comprised of Bryozoa, Brachiopoda and Phoronida (Kocot et al., 2017; Laumer et al., 2019) (Fig. 1a). These animals exhibit “deuterostome-like” features in their development, such as radial cleavage and enterocoely (Zimmer, 1997). We used two brachiopod species, the rhynchonelliform *Terebratalia transversa* and the craniiform *Novocrania anomala*, and one phoronid species, *Phoronopsis harmeri* (Fig. 1b), which show profound differences in mesoderm development such as the time and site of mesoderm emergence, the direction of mesoderm migration and the degree of mesoderm compartmentalization. In particular, in *T. transversa* mesoderm originates at the blastula stage, while the mesoderm of *N. anomala* and *Ph. harmeri* forms at the gastrula stage (*twist* positive cells) (Andrikou et al., 2019; Martín-Durán et al., 2016; Passamaneck et al., 2015). In addition, in *T. transversa* and *Ph. harmeri* mesoderm proliferates in an anterior-to-posterior direction, but in *N. anomala* it follows a posterior-to-anterior direction (Andrikou et al., 2019; Freeman, 1993; Freeman, 2000; Freeman, 2003; Martín-Durán et al., 2016; Nielsen, 1991; Passamaneck et al., 2015; Rattenbury, 1954; Temereva and Malakhov, 2007).

Finally, in *T. transversa* larvae, mesoderm consists of an anterior domain in the apical lobe, an umbrella-like domain in the mantle lobe that projects to four coelomic sacs with chaetae bundles, and a posterior domain in the pedicle lobe (Freeman, 2003; Martín-Durán et al., 2016; Passamaneck et al., 2015; Vellutini and Hejnol, 2016); in *N. anomala* mesoderm compartmentalizes in four pairs of coelomic sacs, with the three posterior ones to project into chaetae bundles (Freeman, 2000; Martín-Durán et al., 2016; Nielsen, 1991; Vellutini and Hejnol, 2016), and in *Ph. harmeri* mesoderm can be distinguished between an anterior domain in the pre-oral lobe with projecting ventro-lateral muscle bands, and a posterior domain that emerges at the larva stage (Andrikou et al., 2019; Rattenbury, 1954; Temereva and Malakhov, 2007) (Figure 1b). We investigated and compared the molecular mechanisms of mesoderm development in these three species, with an emphasis on the role of FGF. Despite our observed differences in the presumptive mesodermal gene regulatory networks (GRN), our results indicate a conservation of the inductive role of FGF signaling pathway in mesoderm with deuterostomes.

## Results

### The spatiotemporal mesodermal patterning differs between *N. anomala, T. transversa* and *Ph. harmeri*

To understand whether the developmental and morphological variations of mesoderm formation between these three species are associated to differences in molecular patterning, we revealed the expression of the transcription factors *twist, mox, six1/2, eya, mef2, dachs, paraxis, foxc, mprx, myod, limpet, foxf* and *nk1* in *N. anomala* and *Ph. harmeri* (Figs 2, 3 S1 and S2). The mesodermal expression of these genes has been previously described in *T. transversa* (Passamaneck et al., 2015). All genes, with the exception of *nk1* (in *N. anomala)* (Fig. S2a), showed mesodermal expression. The earliest mesodermal marker is *twist* (Andrikou et al., 2019; Martín-Durán et al., 2016), whose expression initiates at the early gastrula stage and demarcates the entire mesoderm in both species (Figs 2A2-3 and 3A2-3), indicating that mesoderm originates before its morphological separation from endoderm. *Mox, six1/2* and *eya*, genes commonly involved in mesoderm patterning, are expressed shortly after (Figs 2B2-3, 2C2-3, 2D2-3 and 3B2-3, 3C2-3, 3D2-3). Transcripts of transcription factors often associated with muscle development, such as *myod, limpet* (only in *N. anomala*), *foxf* (Martín-Durán et al., 2016) and the terminal differentiation gene *tropomyosin*, are only detected at the late gastrula stage (Figs 2H4-5, 2I4-5, 2J4-5, 2K4-5 and 3J4-5, 3K4-5) in both organisms, correlating with the formation of musculature (Altenburger and Wanninger, 2010; Temereva and Tsitrin, 2013).

**Fig. 2.**
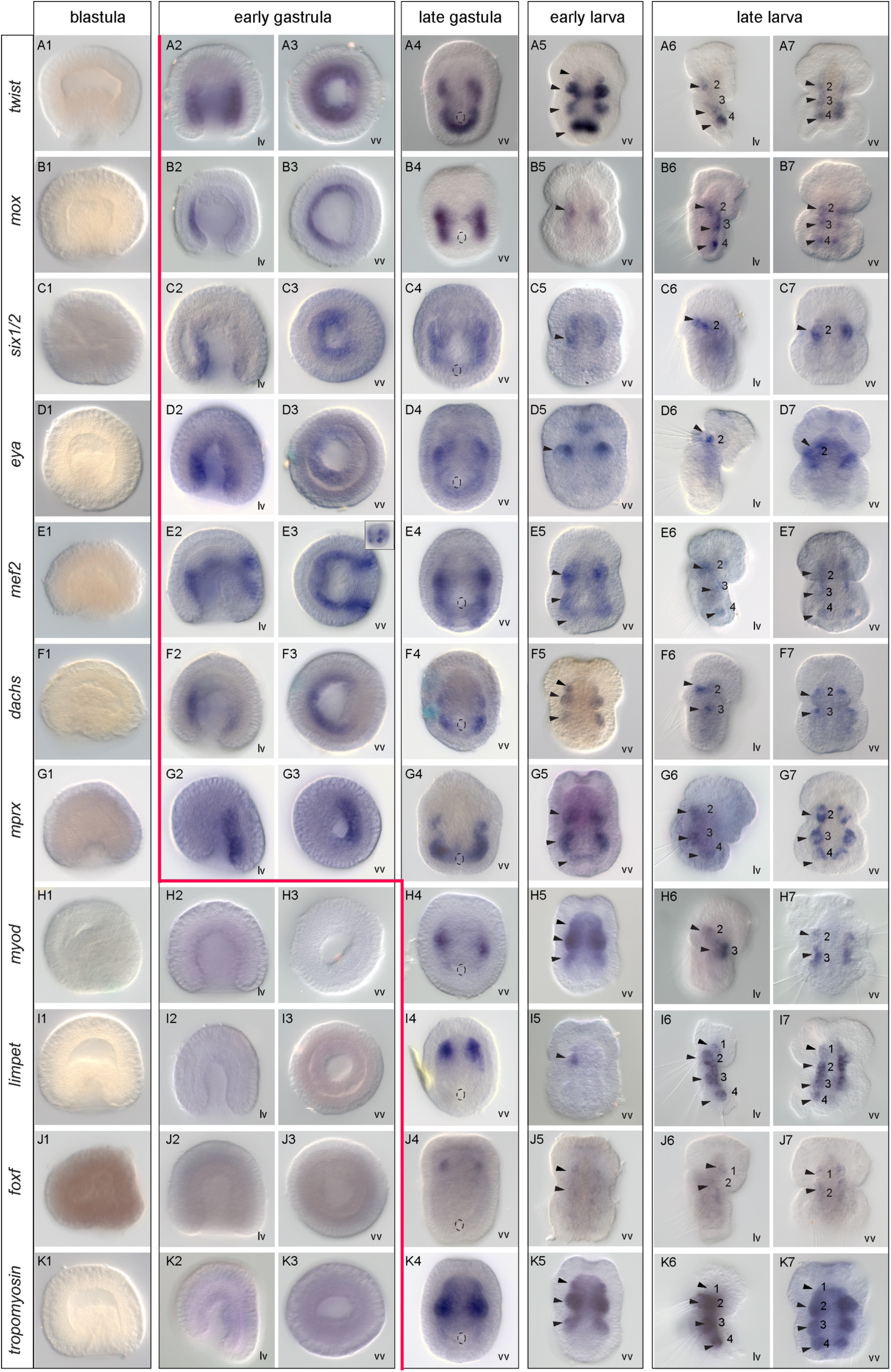
Mesodermal gene expression during *N. anomala* development. WMISH of *twist, mox, six1/2, eya, mef2, dachs, mprx, myod, limpet, foxf* and *tropomyosin* in blastulae, early gastrulae, late gastrulae, early larvae and late larvae of *N. anomala*. The inset in panel E3 shows a different focal plane of the embryo. The position of the blastopore is indicated with a dashed circle in the late gastrula stages. Black arrowheads indicate the coelomic sacs, in which gene expression is detected. The red line marks the onset of mesodermal gene expression. Anterior to the top. lv, lateral view; vv, vegetal view.

**Fig. 3.**
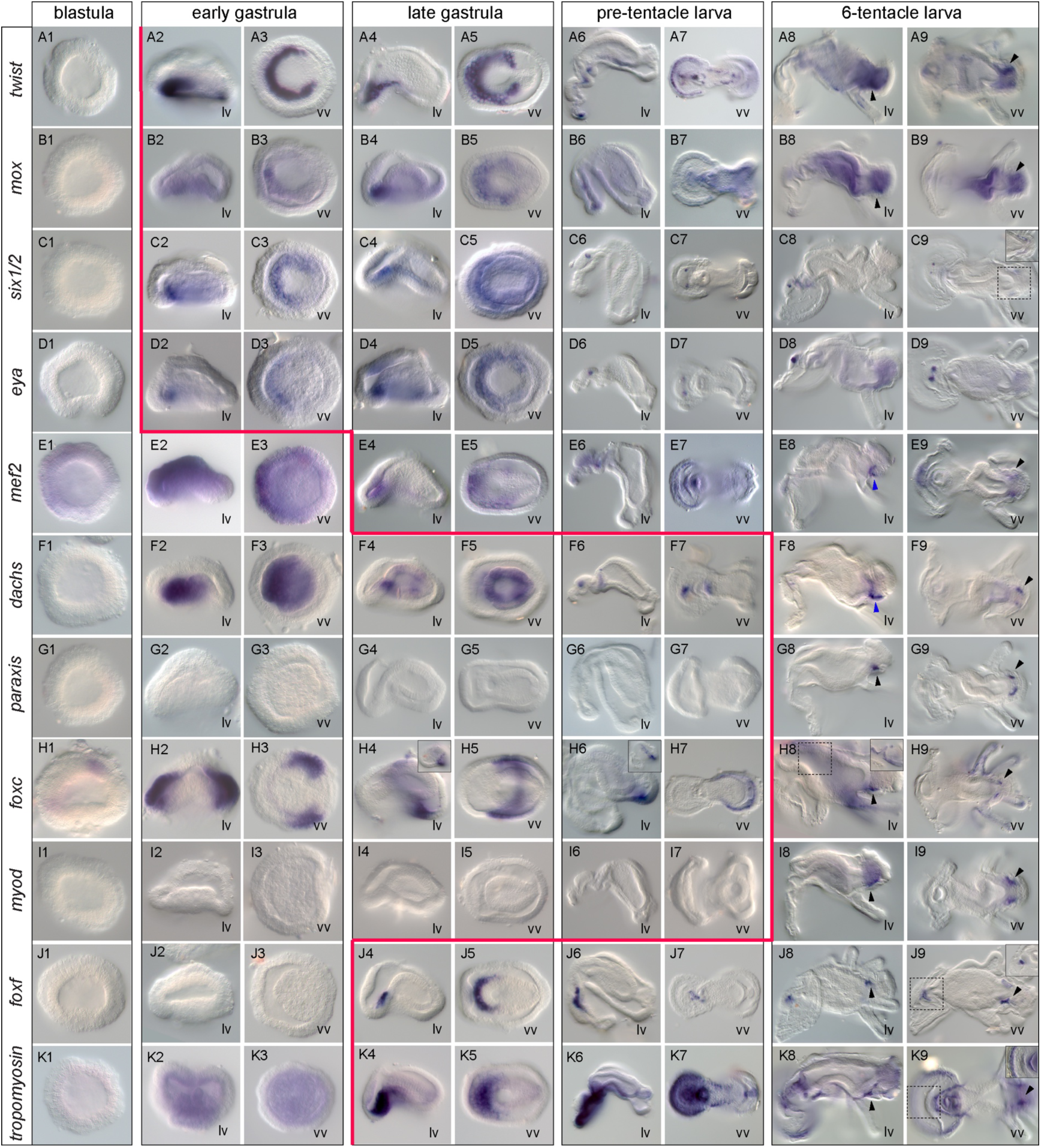
Mesodermal gene expression during *Ph. harmeri* development. WMISH of *twist, mox, six1/2, eya, mef2, dachs, paraxis, foxc, myod, foxf* and *tropomyosin* in blastulae, early gastrulae, late gastrulae, pre-tentacle larvae and 6-tentacle larvae of *Ph. harmeri*. Insets in panels H4 and H6 show different focal planes of the embryos and insets in panels C9, H8, J9 and K9 show higher magnifications of the indicated domains. Black arrowheads indicate expression in the posterior mesoderm. Blue arrowheads show expression in the metasomal sac. The red line marks the onset of mesodermal gene expression. Anterior to the left. lv, lateral view; vv, vegetal view.

However, when comparing these results with data from *T. transversa* (Martín-Durán et al., 2016; Passamaneck et al., 2015), the onset of expression of a number of orthologs varies (Fig. S3). For instance, in *T. transversa* the expression of *mox* and *eya* only starts at the late gastrula stage (Passamaneck et al., 2015), although mesoderm (*twist* positive cells) is already present since the blastula stage (Martín-Durán et al., 2016). Also, *mef2* shows an early mesodermal expression in *N. anomala* (Fig. 2E2-3), but in *T. transversa* (Passamaneck et al., 2015) and *Ph. harmeri* (Fig. 3E4-5) this gene is not activated before the late gastrula stage. A second important difference concerns the spatial patterning of the different subpopulations of mesoderm (Fig. S3). *Twist* is expressed in both anterior and posterior mesodermal subsets in *Ph. harmeri* (Fig. 3A2-9), in *N. anomala* it is expressed in most of the mesoderm but acquires a stronger expression at the posterior one (cs4) (Fig. 2A6-7), and in *T. transversa twist* expression confines in the anterior mesoderm (Passamaneck et al., 2015). The expression of *six1/2* and *myod* is excluded from the posterior (cs4) mesoderm in *N. anomala* (Fig. 2C6-7, H6-7), while in *T. transversa* these genes are expressed in both the anterior (apical) and posterior (pedicle) mesoderm (Passamaneck et al., 2015). In *Ph. harmeri six1/2* and *myod* are restricted in the anterior (Fig. 3C2-5) and posterior mesoderm, respectively (Figs 3I8-9 and S2b). Moreover, *mprx* is solely expressed in the mantle mesoderm of *T. transversa* (Passamaneck et al., 2015), while in *N. anomala* the orthologous gene is expressed in the whole mesoderm (Fig. 2G6-7). *Foxf* and *foxc* expression confines in the anterior mesoderm in both brachiopod species (Passamaneck et al., 2015) (Fig. 2J4-7 and S3) (Martín-Durán et al., 2016), while in *Ph. harmeri foxf* is expressed in both the anterior and posterior mesoderm (Fig. 3J4-9) and transcripts of *foxc* are only detected in the posterior mesoderm (Fig. 3H8-9). *Dachs* seems to be absent from the anterior mesodermal pattering in *Ph. harmeri* (Fig. 3F2-9), while in *T. transversa* it demarcates the entire mesoderm (Passamaneck et al., 2015), and in *N. anomala* it mainly occupies anterior (apical) mesodermal fates (Fig. 2F6-7). Finally, *hox3*, was previously described to be expressed in mid/posterior mesodermal domains in brachiopods (Schiemann et al., 2017), but not in *Ph. harmeri*, where the orthologous gene is solely expressed in the metasomal sac (Gasiorowski and Hejnol, 2020). These data show that in all three organisms the different subsets of mesoderm development exhibit differences not only in the recruitment of regulatory genes but also in their temporal and spatial expression profiles, suggesting diverse mesodermal patterning mechanisms.

### Gene expression of FGF signaling components suggest their possible association with mesoderm and neuronal development

Three FGF receptors were found in *N. anomala* and *T. transversa* but only one in *Ph. harmeri* (Fig. S1). Moreover, all three animals possess one copy of FGF9/16/20 and FGF8/17/18 ligands (Fig. S1). In *T. transversa fgfr1* is expressed in few cells at the vegetal pole at the blastula stage (Fig. 4A1). In gastrulae, transcripts of the gene are demarcating the invaginating endomesoderm (Fig. 4A2) and this expression is retained at the larva stages, in the archenteron, the posterior mesoderm and the developing chaetae sacs (Fig. 4A3). In larvae, *fgfr1* is additionally activated in anterior scattered cells, resembling neurons (Fig. 4A4). *Fgfr2* is strongly expressed in the animal pole of the blastula in a salt and pepper pattern, suggesting neuronal expression (Fig. 4B1). This expression persists in the gastrula and early larva stages (Fig. 4B2-3) but in late larvae it disappears (Fig. 4B4). The third FGF receptor, *fgfr3*, is expressed transiently in blastulae, in few cells of the animal pole (Fig. 4C1), while at the gastrula stage it is also expressed faintly is a small cell cluster of the anterior portion of the invaginating endomesoderm (Fig. 4C2). Larvae are cleared from *fgfr3* transcripts (Fig. 4C3-4). The two ligands also exhibit a very distinct expression from each another. *Fgf9/16/20* is expressed in few cells of the animal pole from the blastula stage up to the larva stage (Fig. 4D1-4). In contrast, *fgf8/17/18* (Vellutini and Hejnol, 2016) starts to be expressed at the blastula stage in an anterior-ventral ectodermal half ring (Fig. 4E1), while in gastrulae, transcripts of the gene are detected in transverse ventral bands reaching the anterior domain of the blastopore, and the future apical organ (Fig. 4E2). In early larvae, *fgf8/17/18* is expressed in two lateral spots, which correspond to the developing chaetae sacs, in anterior cellular patches, as well as in one ventral pair of spots proximal to the mouth and another dorsal pair. Moreover, a new domain of expression at the posterior tip gets activated (Fig. 4E3). Finally, in late larvae, the ventral expression disappears and *fgf8/17/18* is only expressed in the anterior patches, the chaetae sacs and the posterior tip (Fig. 4E4). The analysis of the spatial expression of the three receptors and the two ligands suggests a putative involvement of FGFR1 and FGF8/17/18 in mesoderm development (see co-expression of *fgfr1* and *fgf8/17/18* in Fig. S4).

**Fig. 4.**
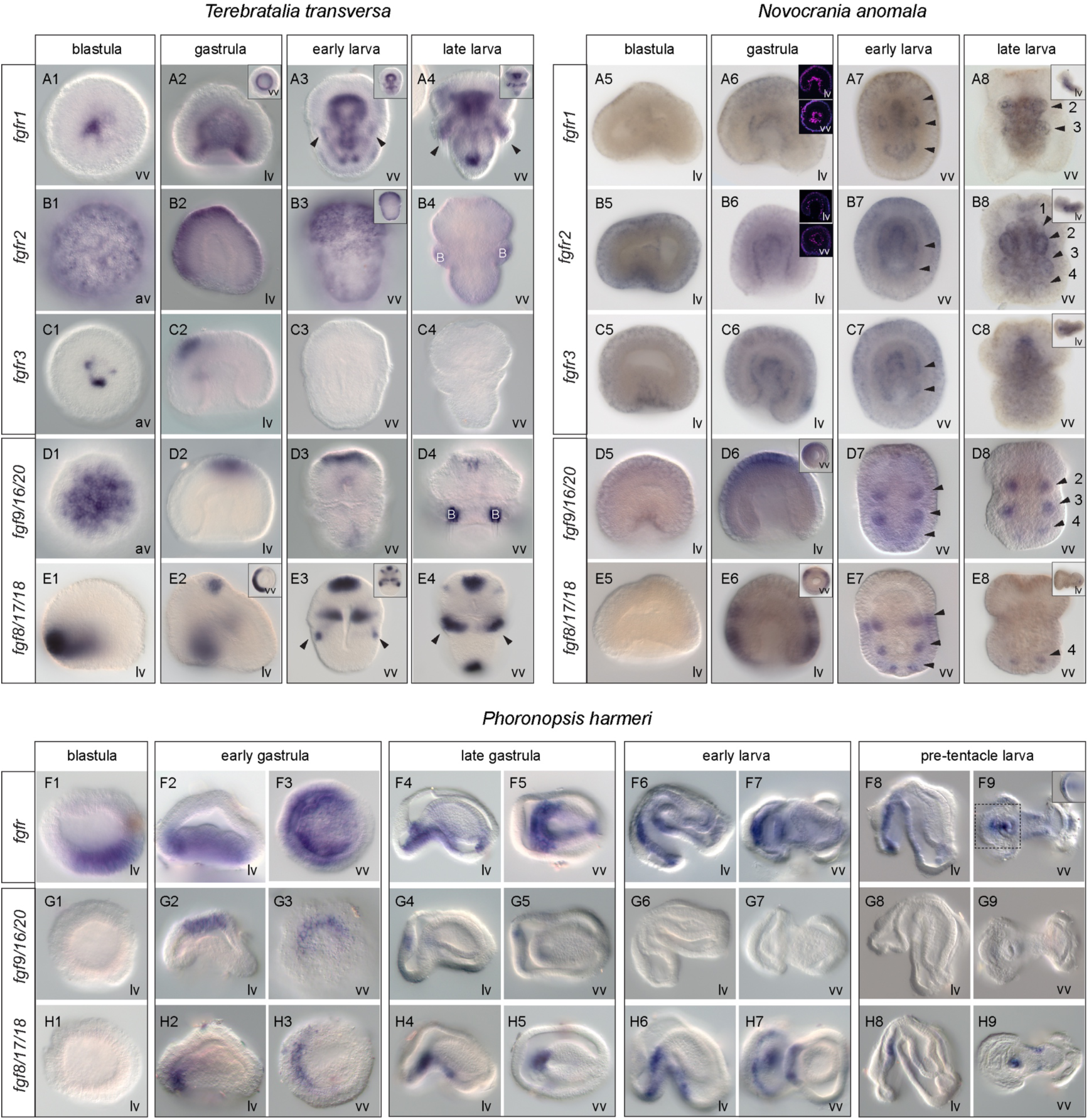
Gene expression of FGF signaling components in lophophorates. WMISH of *fgfr1, fgfr2, fgfr3, fgf8/11/18* and *fgf9/16/20* during the blastula, gastrula, early larva and larva stages of development of *T. transversa, N. anomala* and *Ph. harmeri*. Insets in panels A3, A4, B3 and E3 show different focal planes of the embryos, insets in panels A2, E2, D6 and E6 show vegetal views, panel B2 shows an animal view and panels A8, B8, C8, D8, E8 shows lateral views. Insets in A6 and B6 show fluorescent WMISH. The inset in panel A9 shows a higher magnification of the indicated domain. Black arrowheads indicate the coelomic sacs, in which gene expression is detected. For *T. transversa, N. anomala* anterior to the top and for *Ph. harmeri* anterior to the left. av, animal view; lv, lateral view; vv, vegetal view.

In *N. anomala* none of the FGF signaling components is expressed at the blastula stage, which differs from what is observed in *T. transversa* (Fig. 4A5, B5, C5). The expression of all *fgf* receptors is detected at the gastrula stage, in the invaginating archenteron and the developing coelomic sacs (Fig. 4A6, B6, C6). Additionally, transcripts of *fgfr1* are found in the anterior ectoderm (Fig. 4A6). In early larvae, *fgfr1* is expressed in anterior ectodermal cells, the invaginating mesoderm and the tip of the archenteron (Fig. 4A7). At the late larva stage, *fgfr1* expression confines in the two anterior pairs (cs2-cs3) of chaetae sacs (Fig. 4A8). *Fgfr2* and *fgfr3* are mainly expressed in the forming archenteron and the invaginating mesoderm at the early larva stage (Fig. 4B7, C7) but *fgfr2* is further detected in two anterior-lateral ectodermal patches (Fig. 4B7). Finally, in larvae, *fgfr2* is expressed in all four pairs (cs1-cs2-cs3-cs4) of coelomic sacs (Fig. 4B8), while *fgfr3* is only expressed in the mouth region (Fig. 4C8). The two ligands also start to be expressed during gastrulation. *Fgf9/16/20* expression is initially detected in the anterior ectoderm (Fig. 4D6) but in larvae the ectodermal expression fades and a new mesodermal domain appears in the three coelomic pairs (cs2-cs3-cs4) of chaetae sacs (Fig. 4D7-8). *Fgf8/17/18* (Vellutini and Hejnol, 2016) is expressed in two ectodermal bands that encircle the gastrula, one more posterior near the blastopore and another at the mid part of the embryo (Fig. 4E6). In early larvae, transcripts of the gene are detected in the three developing pairs of chaetae sacs (cs2-cs3-cs4) and the ectodermal patches adjacent to the first pair (cs2) (Fig. 4E7) and at the late larva stage the expression of *fgf8/17/18* is restricted to the most posterior pair (cs4) of chaetae sacs (Fig. 4E8). Based on their expression, these data suggest that all three receptors and both ligands are possibly related to mesoderm development in *N. anomala*.

In *Ph. harmeri*, the only FGF receptor is already expressed at the blastula stage, in the vegetal pole (Fig. 4F1). At the early gastrula stage, the gene is expressed in an anterior ventro-lateral cell population of the vegetal plate, the presumptive mesoderm, the anterior blastoporal lip, as well as the anterior ectoderm that will give rise to the apical organ (Fig. 4F2-3). In late gastrulae, the gene is expressed in anterior migrating mesodermal cells and a posterior cell cluster located adjacent to the developing intestine (Fig. 4F4-5). At the early larva and pre-tentacle larva stages, the expression of *fgfr* remains in clusters of cells of the pre-oral mesoderm, two ventro-lateral mesodermal tiers, the posterior cell cluster, the ventral ectoderm and the apical organ (Fig. 4F6-9). *Fgf9/16/20* and *fgf8/17/18* exhibit very different expression profiles. *Fgf9/16/20* is transiently expressed in the anterior mesoderm and the forming apical organ until the late gastrula stage (Fig. 4G2-5). *Fgf8/17/18* exhibits a more dynamic expression, detected in the anterior lip of the blastopore, the anterior-ventral ectoderm and the anterior endoderm in early gastrulae (Fig. 4H2-3), while in later gastrulae and larvae *fgf8/17/18* is expressed in the anterior-ventral ectoderm of the oral hood, a postero-ventral group of ectodermal cells and the mouth (Fig. 4H4-9). These data show that also in *Ph. harmeri*, the expression of FGFR and FGF8/17/18 is possibly associated to mesoderm formation (see coexpression of *fgfr* and *fgf8/17/18* in Fig. S4). Overall, FGF signaling is likely involved in mesoderm development, as well as neuroectodermal patterning and apical organ formation in all three organisms. A summary of the expression of the FGF signaling components in *N. anomala, T. transversa* and *Ph. harmeri* is provided in Fig. S5.

### Perturbation of FGF signaling results in failure in mesoderm formation in *N. anomala* but not in *T. transversa*

Based on the mesoderm-related expression of some FGF signaling components we hypothesized that FGF might be involved in mesoderm development in brachiopods. To test this, we treated embryos at different developmental stages with SU5402, a selective inhibitor of FGFR (Mohammadi et al., 1997) (for a summary of treatments see Fig. S6). SU5402 treatment abolished the formation of chaetae sacs and neuropile in larvae of both brachiopod species (Fig. 5). However, a difference was observed in mesoderm formation.

**Fig. 5.**
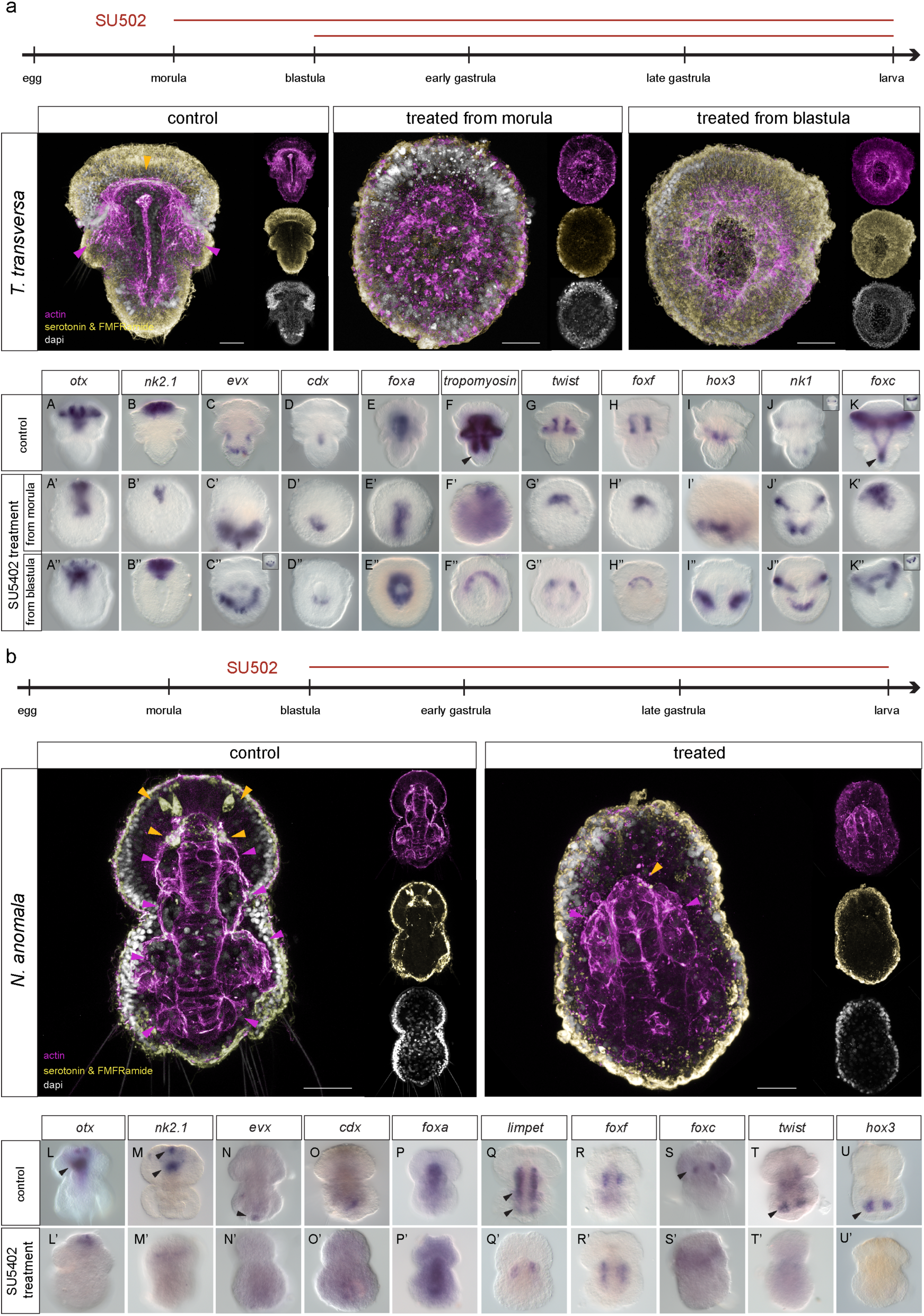
SU5402 treatments in brachiopods. (a) Immunohistochemistry of markers of the nervous system (serotonin, FMFRamide) and musculature (actin) in *T. transversa* morula and blastula embryos treated with 20 uM SU5402 and fixed at larva stage. WMISH of anterior (*otx, nk2.1*), posterior *(evx*) and endodermal *(foxa, cdx)* genes, musculature (*tropomyosin)*, anterior mesoderm (*twist, foxf)*, mid mesoderm (*hox3*) and posterior mesoderm *(nk1, foxc)* in *T. transversa* morula and blastula embryos treated with 20 uM SU5402 and fixed at larva stage. Insets in panels J, K, C’’ and K’’ show different focal planes of the embryos. (b) Immunohistochemistry of markers of the nervous system (serotonin, FMFRamide) and musculature (actin) in *N. anomala* blastula embryos treated with 20 uM SU5402 and fixed at larva stage. WMISH of anterior (*otx, nk2.1*), posterior genes (*evx*) and endodermal *(foxa, cdx)* genes, the whole mesoderm (*limpet)*, anterior mesoderm (*foxf, foxc)* and posterior mesoderm *(twist, hox3)*, in *N. anomala* blastula embryos treated with 20 uM SU5402 and fixed at larva stage. Black arrowheads indicate the domains of expression that are absent in the treated embryos. Yellow arrowheads indicate the neuropile and magenta arrowheads the musculature associated to coelomic and chaetae sacs. All panels depict embryos in vegetal view. Every fluorescent image is a full projection of merged confocal stacks and nuclei are stained with DAPI. Anterior to the top. Drawings are not up to scale. Scale bar: 20 um.

*T. transversa* treated larvae were not compartmentalized in apical, mantle and pedicle lobes, but instead were remaining spherical with an open blastopore, when we treated them from the blastula stage (Fig. 5a). The chaetae bundles were not formed, and the musculature was impaired (Fig. 5a). To ensure that we inhibit FGFR before mesoderm originates, we also treated larvae from the morula stage, which resulted in spherical embryos without a blastopore (Fig. 5a). To understand whether this truncated phenotype was due to a failure of axial elongation or a disruption of the anterior-posterior patterning, we tested the expression of anterior (*otx, nk2.1*) (Fig. 5aA-A’’, B-B’’) and posterior markers (*evx*) (Martín-Durán et al., 2016) (Fig. 5aC-C’’) and found them unaffected. The expression of the endoderm markers *cdx* and *foxa* (Martín-Durán et al., 2016) was also unaltered (Fig. 5aD-D’’, E-E’’). We then examined the treated animals for the muscle differentiation marker *tropomyosin* (Passamaneck et al., 2015) and a loss of posterior expression was observed (Fig. 5aF-F’’). We then tested the expression of the transcription factors *twist* and *foxf*, markers of anterior/apical mesoderm (Passamaneck et al., 2015) (Fig. 5aG-G’’, H-H’’), *hox3*, a marker of mid/mantle mesoderm (Schiemann et al., 2017) (Fig. 5aI-I’’), and *nk1*, a marker of posterior/pedicle mesoderm (Passamaneck et al., 2015) (Fig. 5aJ-J’’) and found them unchanged. Interestingly, the expression of *foxc*, a marker of the most posterior mesoderm (Passamaneck et al., 2015) was lost (Fig. 5aK-K’’), suggesting a role of this gene in a later, differentiation step of mesoderm (see also Fig. S7). Moreover, when we treated animals from a later developmental stage (mid gastrula), the phenotype was milder, the posterior mesodermal expression of *foxc* was recovered and *tropomyosin* expression was extended more posteriorly compared to the larvae treated from the blastula stage (Fig. S8a). These results suggest that in *T. transversa*, FGF signaling is involved in neuropile formation, coordination of morphogenetic movements of gastrulation and axial elongation. It does not have a role in mesoderm induction (see also Fig. S9), but instead in mesoderm migration and differentiation (as seen from the loss of chaetae sacs). Since in *T. transversa* the direction of axial elongation is taking place from anterior-dorsal to posterior (Freeman, 1993; Martín-Durán et al., 2016), the inhibition in posterior axial elongation is probably coupled with the failure in mesoderm migration and differentiation.

*N. anomala* treated animals did not exhibit the same phenotype that we observed in *T. transversa* (Fig. 5b). Larvae treated from the blastula stage were smaller than the controls, but they were not spherical, and they possessed an elongated archenteron without a mouth opening (Fig. 5b). Also, mesoderm was impaired; only one out of the four pairs of coelomic sacs was present and none of the chaetae bundles were formed (Fig. 5b). When we looked at the anterior-posterior patterning genes (Martín-Durán et al., 2016) we observed an apical reduction and a complete loss of expression in the mouth region of the genes *otx* and *nk2.1* (Fig. 5bL-L’, M-M’). The most posterior fate was also impaired, as shown from the reduced expression of *evx* (Fig. 5bN-N’). The expression of the endodermal markers *cdx* and *foxa* (Martín-Durán et al., 2016), however, remained unaffected (Fig. 5bO-O’, P-P’). We then tested the expression of *limpet*, a pan-mesodermal differentiation marker in this species. We saw a severe reduction of expression and detection in only one anterior coelomic sac (Fig. 5bQ-Q’). The expression of the anterior mesodermal marker *foxf* (Martín-Durán et al., 2016) was not affected (Fig. 5bR-R’), but instead transcripts of *foxc* (Martín-Durán et al., 2016), which in control larvae confine in the most anterior coelomic sac (cs1), were lost (Fig. 5bS-S’), suggesting that the formation of cs1 was abolished. Finally, the expression of the posterior mesodermal markers *twist* (Martín-Durán et al., 2016) and *hox3* (Schiemann et al., 2017) were inhibited (Fig. 5bT-T’, U-U’). Similar results were obtained when the embryos were treated from the gastrula stage, with the exception of *hox3*, the expression of which was partly recovered (Fig. S8b), indicating that the input of FGF on *hox3* occurs sometime between the blastula and gastrula stage. In order to understand which step of mesoderm development had been compromised, we also tested the expression of the mesodermal marker *twist* in gastrulae treated from the blastula stage on, and found it downregulated, suggesting that mesoderm induction was compromised (Fig. S9). These data suggest that in *N. anomala*, FGF is involved in anteroposterior patterning, neuropile formation, mouth formation, as well as the most anterior and posterior mesoderm formation and mesoderm differentiation (as seen from the loss of chaetae sacs). However, an impact in axial elongation is not evident, as shown in *T. transversa*, further supported by the fact that in *N. anomala* the direction of axial elongation is occurring from posterior-ventral to anterior (Freeman, 2000; Martín-Durán et al., 2016; Nielsen, 1991).

### FGF signaling is upstream of mesoderm induction in *Ph. harmeri*

To test whether the role of FGF signaling in mesoderm induction extends beyond the brachiopod lineage, we also treated embryos of *Ph. harmeri* with SU5402 at different developmental stages (for a summary of treatments see Fig. S6). SU5402 treatment from the blastula stage inhibited the formation of musculature and the apical organ of pre-tentacle larvae (Fig. 6a). The treated larvae exhibited a truncated phenotype with a shorter archenteron that didn’t have a visible opening (Fig. 6a). The expression of the mesodermal markers *twist, six3/6* (Andrikou et al., 2019) and *foxf* was abolished (Fig. 6aA-F’), and the same was observed for the markers of the apical organ *six3/6* and *otx* (Andrikou et al., 2019) (Fig. 6aC-D’, I-J’). The ectodermal expression of *foxc*, was only dorsally affected (Fig. 6aK-L’). Finally, the posterior endodermal expression of *nk2.1* (Fig. 6aG-H’) and the expression of *foxa* (Andrikou et al., 2019) in the mouth (Fig. 6aM-N’) were downregulated. A treatment from the gastrula stage on, resulted in a milder phenotype (Fig. S10). Some mesodermal cells were present, the apical organ was recovered, and the mouth was formed (Fig. S10). Moreover, the posterior endodermal expression of *nk2.1* -as well as *foxa* expression in the mouth-was unaffected (Fig. S10). To test whether this was due to a failure in mesoderm induction, we also tested the expression of the mesodermal markers *twist* and *six3/6* in gastrulae treated from the blastula stage on, and found it abolished (Fig. S9). These results show that FGF signaling is upstream of anteroposterior patterning, apical organ formation, gastrulation movements and mesoderm formation in *Ph. harmeri*. Moreover, an additional participation of FGF in mesoderm migration is also possible due to the coexpression of the mesodermal marker *twist* and *fgfr* throughout the development (Fig. 6b).

**Fig. 6.**
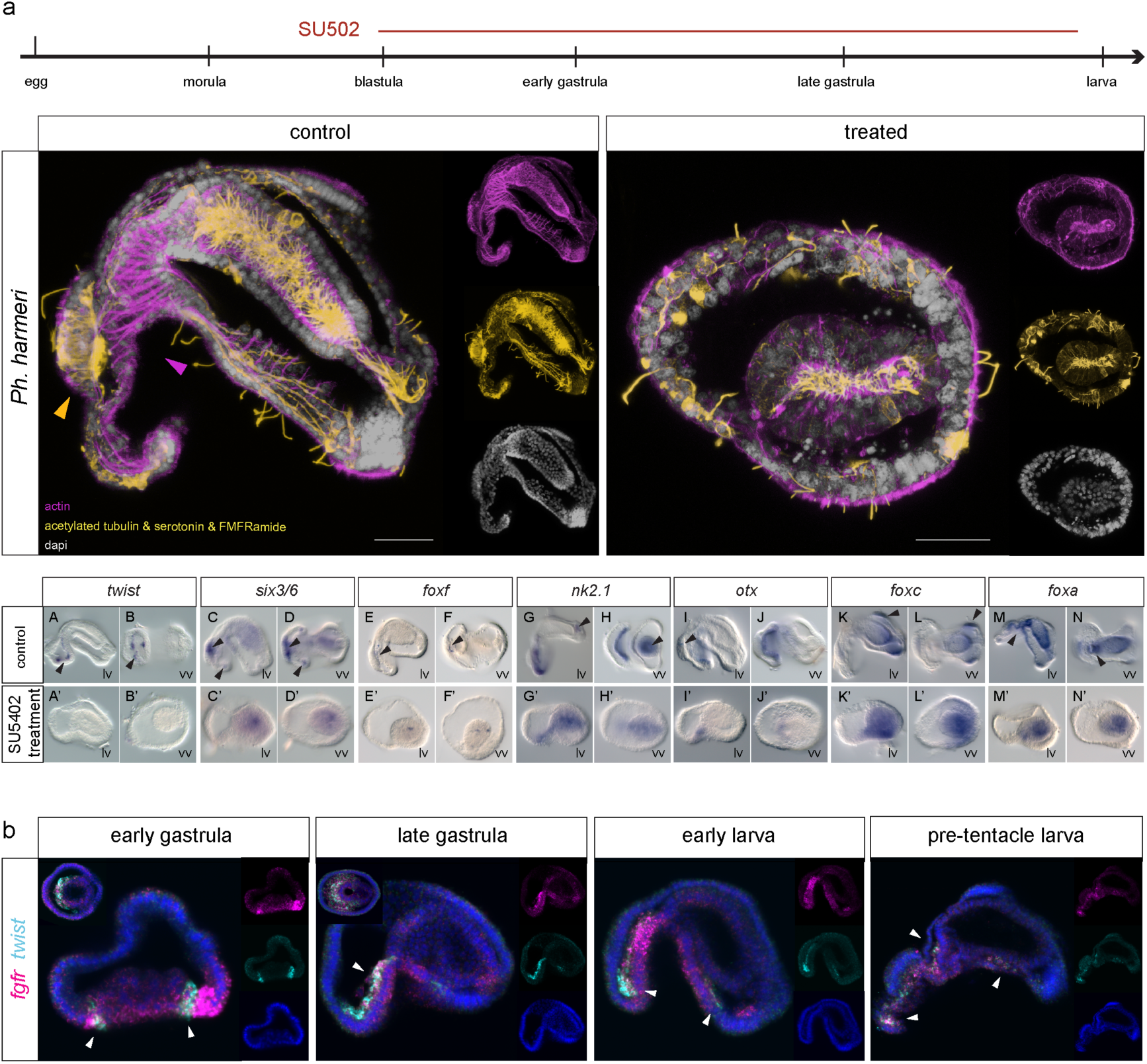
SU5402 treatments in *Ph. harmeri*. (A) Immunohistochemistry of markers of the nervous system (acetylated tubulin, serotonin, FMFRamide) and musculature (actin) in *Ph. harmeri* blastula embryos treated with 20 uM SU5402 and fixed at larva stage. Yellow arrowheads indicate the apical organ and magenta arrowheads the esophageal musculature. WMISH of mesodermal (*twist, six3/6, foxf)*, anterior (*six3/6, otx, nk2.1*), postero-ventral (*foxc*) and endodermal (*foxa, nk2.1*) genes in *Ph. harmeri* blastula embryos treated with 20 uM SU5402 and fixed at larva stage. Black arrowheads indicate the domains of expression that are absent in the treated embryos. (B) Co-expression analysis of *fgfr* (magenta) and *twist* (cyan) by double fluorescent WMISH in gastrula and larva stages of development of *Ph. harmeri*. Right insets show embryos in vegetal view. White arrowheads indicate co-expression. Every fluorescent image is a full projection of merged confocal stacks and nuclei are stained with DAPI. Anterior to the left. lv, lateral view; vv, vegetal view. Scale bar: 20 um.

Overall, our data suggest an evident role of FGF signaling in mesoderm development in the lineage of lophophorates. Moreover, a conserved involvement of FGF in anteroposterior patterning, neuron formation, morphogenetic movements of gastrulation and axial elongation is witnessed.

## Discussion

### Expression dynamics of the mesodermal gene battery

Nearly all the genes we studied are expressed during mesodermal development in the investigated lophophorate species however, the temporal expression dynamics and spatial recruitment of some genes differ (Fig. S3). While *twist, six1/2* and *foxf* are expressed in a similar sequential manner in all three organisms, the remaining genes occupy different temporal regulatory positions. The spatial utilization of the genetic repertoire in the differentiated subsets of mesoderm exhibits only few cases of shared spatial similarity in all three species (e.g. *mef2*, which demarcates the entire mesoderm, and *eya*, which is mostly expressed in the anterior mesoderm), but the other genes show differences in their spatial transcript distribution. Overall, these results suggest that mesoderm development in lophophorates utilizes a similar set of transcription factors, but their hierarchical deployment differs, suggesting profound differences in their mesodermal patterning and mesoderm regionalization. Data from bryozoans, the potential sister group of phoronids, suggest similar spatial differences in the mesodermal patterning, such as the posterior expression of *foxc* (Vellutini et al., 2017). Moreover, comparative studies of the expression profiles of endomesoderm and ectomesoderm in lophotrochozoans have revealed some intriguing differences, such as the confinement of *twist* expression in the ectomesoderm of the mollusks *Crepidula fornicata* (Perry et al., 2015), *Patella vulgata* (Nederbragt et al., 2002) and the annelid *Capitella teleta* (Dill et al., 2007) but not in the annelids *A. virens* and *P. dumerilii*, where *twist* is expressed in both sources of mesoderm (Kozin et al., 2016; Pfeifer et al., 2014; Steinmetz, 2006).

It thus becomes evident that the spatial and temporal differences in lophophorate mesoderm development are observed in more spiralian taxa, which indicates a diversification of mesodermal developmental programs and their underlying GRNs. Different circuitries of GRNs orchestrating the formation of homologous mesodermal derivatives have been described in some animals and support the idea that the evolution of GRNs is mainly based on the developmental regulatory demands of each network (Andrikou and Arnone, 2015; Erkenbrack, 2016; Erkenbrack et al., 2018; Hinman and Davidson, 2007). Therefore, alterations in GRN circuitries do not necessarily reflect convergent evolution of the resulting tissues (Davidson and Erwin, 2006; Peter, 2020), but can also be a product of developmental system drift (True and Haag, 2001).

### FGF signaling upstream of different mesodermal populations

FGF signaling is required for the formation of all or most mesoderm e.g. in hemichordates (Fan et al., 2018; Green et al., 2013) and tunicates (Davidson et al., 2006; Imai et al., 2002; Kim and Nishida, 2001; Yasuo and Hudson, 2007), or a subset of mesoderm e.g. in vertebrates (Amaya et al., 1993; Draper et al., 2003; Fletcher et al., 2006; Fletcher and Harland, 2008; Yamaguchi et al., 1994), cephalochordates (Bertrand et al., 2011), sea urchins (Andrikou et al., 2015) and nematodes (Photos et al., 2006) (S11). According to our results, this is similar to lophophorates, where FGF acts on different levels of mesoderm development. While *N. anomala* utilizes FGF to form two subsets of mesoderm, in *Ph. harmeri* FGF is upstream of the formation of all mesoderm. It remains unclear why mesodermal subpopulations differ in their promoting requirements and deploy different signals. The acquisition of different signaling pathways, with distinct spatiotemporal expression dynamics and inductive properties, can act as a relay mechanism of the initial signal but can also exhibit diverse functions. An example is the recruitment of Nodal in vertebrate development, which although it interacts synergistically with FGF in promoting mesoderm, it also acts differentially in the induction of mesodermal populations (Kimelman, 2006; Mathieu et al., 2004).

### Implications of mesoderm development in gastrulation and axial elongation

Besides having a role in mesoderm development, FGF signaling has conserved functions in neural development and morphogenetic movements of gastrulation in an array of investigated organisms (Fig. S11). In deuterostomes, FGF is involved both in gastrulation (Amaya et al., 1991; Bertrand et al., 2011; Röttinger et al., 2008) and neural induction (Bertrand et al., 2003; De Robertis and Kuroda, 2004; Garner et al., 2016). Also, in the two well-studied ecdysozoans *D. melanogaster* and *C. elegans*, FGF signaling is upstream of axon guidance (Bülow et al., 2004; García-Alonso et al., 2000), and cell migration during gastrulation (in *D. melanogaster*) (Leptin and Affolter, 2004). In the remaining protostomes data are limited to gastropods and Platyhelminthes, where FGF signaling is involved in neural development (Cebrià et al., 2002; Pollak et al., 2014). Most likely, the involvement of FGF in these developmental processes was already present before the cnidarian-bilaterian split, as witnessed in sea anemones, where FGF seem to act upon gastrulation (Matus et al., 2007), neural development (Matus et al., 2007) and is upstream of apical organ formation (Rentzsch et al., 2008). Our study revealed similar roles of FGF signaling in the investigated lophophorate species. In particular, all three species exhibited defects in their apical organ/neuropile formation, as well as loss of a number of differentiated neurons (e.g. serotonergic neurons in Fig. 5b). Moreover, they all showed impaired gastrulation to some degree. Impaired gastrulation can occur either as a direct or indirect effect mediated by a failure in mesoderm formation. For example, in *T. transversa*, where mesoderm is formed independently of FGF signaling, most likely the role of FGF is only morphogenetic, in orchestrating cell movements during gastrulation. However, in the other two species, where FGF is involved in mesoderm formation, it is still unclear whether the observed failure in gastrulation movements after FGF inhibition is direct, or indirect caused by the lack of mesoderm formation.

Another outcome of this study concerns the evident relationship witnessed between mesoderm development and posterior axis elongation in *T. transversa*. The expression of *fgf8/11/18* mRNA in the growing posterior tip of the embryo in relation to *fgfr1* (Fig. S4), suggests that FGF8/11/18 might act as a gradient that progressively coordinates the posterior elongation of the embryo and mesoderm differentiation, similar to what has been described in vertebrates (Dubrulle and Pourquie, 2004). The role of FGF signaling in the posterior axial elongation of *T. transversa* resembles mechanisms typically deployed by a number of deuterostomes in order to coordinate embryonic patterning and elongation of posterior tissues: in the tunicate species *Ciona intestinalis*, FGF, together with the canonical WNT and Retinoic Acid (RA) signaling, are establishing the posterior patterning of the tail (Pasini et al., 2012), in amphioxus FGF and RA are coordinating the posterior elongation (Bertrand et al., 2015), and in vertebrates the AP axial elongation of the trunk is mediated by an antagonistic interplay of FGF and RA, where high RA or low FGF results in cessation of posterior embryonic regions (Deng et al., 1994; Partanen et al., 1998; Yamaguchi et al., 1994). Data from RA signaling are currently unavailable in lophophorates; however, WNT signaling also appears to be involved in axis patterning and the establishment of posterior identities (Martín-Durán et al., 2016; Vellutini and Hejnol, 2016), which suggest a conserved coordinated function of WNT and FGF signaling in regulating posterior axial elongation.

To summarize, the data provide support for a conserved involvement of FGF signaling in gastrulation movements and axial elongation, with the phenotypic severity to vary, depending on the developmental mode of mesoderm formation of the investigated species. A coordinated interplay of WNT and FGF signaling in regulating posterior axial elongation seems also likely, suggesting a putative conserved developmental signaling mechanism in orchestrating posterior regionalization.

### The recurrent use of FGF signaling in mesoderm formation

The role of FGF signaling in mesoderm induction was thought to be restricted to deuterostomes. After investigating three species of lophophorates, we are able to show that the mesoderm-inducing ability of this pathway extends beyond the lineage of deuterostomes. However, signaling pathways are often deployed as upstream “plug-in” devices and can be co-opted and exchanged to serve different developmental processes within and among species (Davidson and Erwin, 2006). To determine whether the involvement of FGF signaling in mesoderm formation was already present in the last common ancestor of Bilateria, or whether it was independently co-opted in the lineage of lophophorates, functional studies from more spiralian taxa are required. So far, the only available data in favour of a putative conserved role of FGF in mesoderm induction come from studies in mollusks, where MAPK, one of the major downstream targets of FGF signaling (Ornitz and Itoh, 2015), is upstream of endomesoderm specification (Koop et al., 2007; Kozin et al., 2013; Lambert, 2008; Lambert and Nagy, 2001; Lambert and Nagy, 2003). Moreover, a putative conserved FGF signaling event on mesoderm induction is possibly also present in bryozoans, as suggested from the activation of the MAPK pathway in the mesodermal precursor 3D (Vellutini et al., 2017), which would imply that the inductive role of FGF signaling in mesoderm was present at least in the last common ancestor of lophophorates.

## Materials and methods

### Animal systems

Gravid adult specimens were collected in Bodega bay, California, USA (*Phoronopsis harmeri* Pixell, 1912), in Friday Harbor Laboratories, U.S.A. (*Terebratalia transversa* Sowerby, 1846), in Espeland Marine Biological Station, Norway (*Novocrania anomala* Müller, 1776) and spawned as previously described (Freeman, 1993; Freeman, 2000; Rattenbury, 1954). The embryos were kept in clean seawater and collected at various stages of development.

### Gene cloning and orthology assignment

Putative orthologous sequences of genes of interest were identified by tBLASTx search against the transcriptomes of *Terebratalia transversa, Novocrania anomala* and *Phoronopsis harmeri*. Gene orthology of genes of interest identified by tBLASTx was tested by reciprocal BLAST against NCBI Genbank and followed by phylogenetic analyses. Amino acid alignments were made with MUSCLE. IQ-tree (version 2.0.5) was used to conduct a maximum likelihood phylogenetic analysis. Fragments of the genes of interest were amplified from cDNA of *T. transversa, N. anomala* and *Ph. harmeri* by PCR using gene specific primers. PCR products were purified and cloned into a pGEM-T Easy vector (Promega, USA) according to the manufacturer’ s instruction and the identity of inserts confirmed by sequencing.

### SU5402 treatments

SU5402 was dissolved in DMSO to a final concentration of 5 um, 10 uM and 20 μM. Higher concentrations than these were lethal to the embryos. SU5402 was added at morula, blastula and gastrula stages up to the fixation stage. A corresponding volume of DMSO was added in the control embryos. Solutions were changed every 24 h. A table summarizing the drug treatments and observed phenotypes is seen in Fig. S6.

### Whole Mount *In Situ* Hybridization

Embryos were manually collected, fixed in 4% paraformaldehyde in SW for 60 minutes, permeabilised in 100% Methanol overnight and processed for *in situ* hybridization as described in (Andrikou et al., 2019; Martín-Durán et al., 2016; Santagata et al., 2012). Labeled antisense RNA probes were transcribed from linearized DNA using digoxigenin-11-UTP (Roche, USA) according to the manufacturer’s instructions.

### Whole Mount Immunohistochemistry

Embryos were permeabilised in 100% Methanol for 1 hour, digested with Proteinase K (10 µg ml^−1^) for 5 minutes, fixed in 4% paraformaldehyde in SW for 30 minutes, washed for 3 hours in 1% PTX, washed in PBT and incubated in 4% sheep serum in PBT for 30 min. The animals were then incubated with commercially available primary antibodies (anti-acetylated tubulin mouse monoclonal antibody (Sigma-Aldrich); anti-actin mouse monoclonal antibody (Seven Hills Bioreagents); anti-serotonin, rabbit monoclonal antibody (Sigma-Aldrich); anti-FMFRamide rabbit monoclonal antibody (Immnunostar); anti-HB9 rabbit monoclonal antibody (Invitrogen) overnight at 4°C, washed 3 times in PBT, and followed by incubation in 4% sheep serum in PBT for 30 min. Specimens were then incubated with secondary anti-rabbit and anti-mouse antibodies Alexa Fluor overnight at 4°C followed by 3 washes in PTW. Nuclei were stained with DAPI (Molecular probes) and F-actin was visualized with BODIPY FL Phallacidin (Molecular probes).

### Documentation

Colorimetric WMISH specimens were imaged with a Zeiss AxioCam HRc mounted on a Zeiss Axioscope A1 equipped with Nomarski optics and processed through Photoshop CS6 (Adobe). Fluorescent-labeled specimens were analyzed with a SP5 confocal laser microscope (Leica, Germany) and processed by the ImageJ software version 2.0.0-rc-42/1.50d (Wayne Rasband, NIH). Fig. plates were arranged with Illustrator CS6 (Adobe).

## Acknowledgements

We thank all members from Hejnol’s group for contributing with animal collection and spawning. We are grateful to Chris Lowe for hosting us in his lab for *Phoronopsis harmeri* collection and spawning. We also thank Paul Bump from Lowe lab and Karl Menard from Bodega Bay Marine Lab for their help with collecting *Phoronopsis harmeri* adults. We are grateful to the “Centennial” boat crew of Friday Harbor and the Espeland marine biological station personnel for the *T. transversa* and *N. anomala* collections, respectively.

## Authors’ contributions

CA designed the study, performed the collections and conducted the experiments. CA and AH performed the collections, analyzed the data and wrote the manuscript.

## Data availability

All newly determined sequences have been deposited in GenBank under the accession numbers xxxxx. Primer sequences are available on request.

