## Supplementary information for "FGF signaling induces mesoderm in members of Spiralia"

**Index**

Supplementary figures 1-12

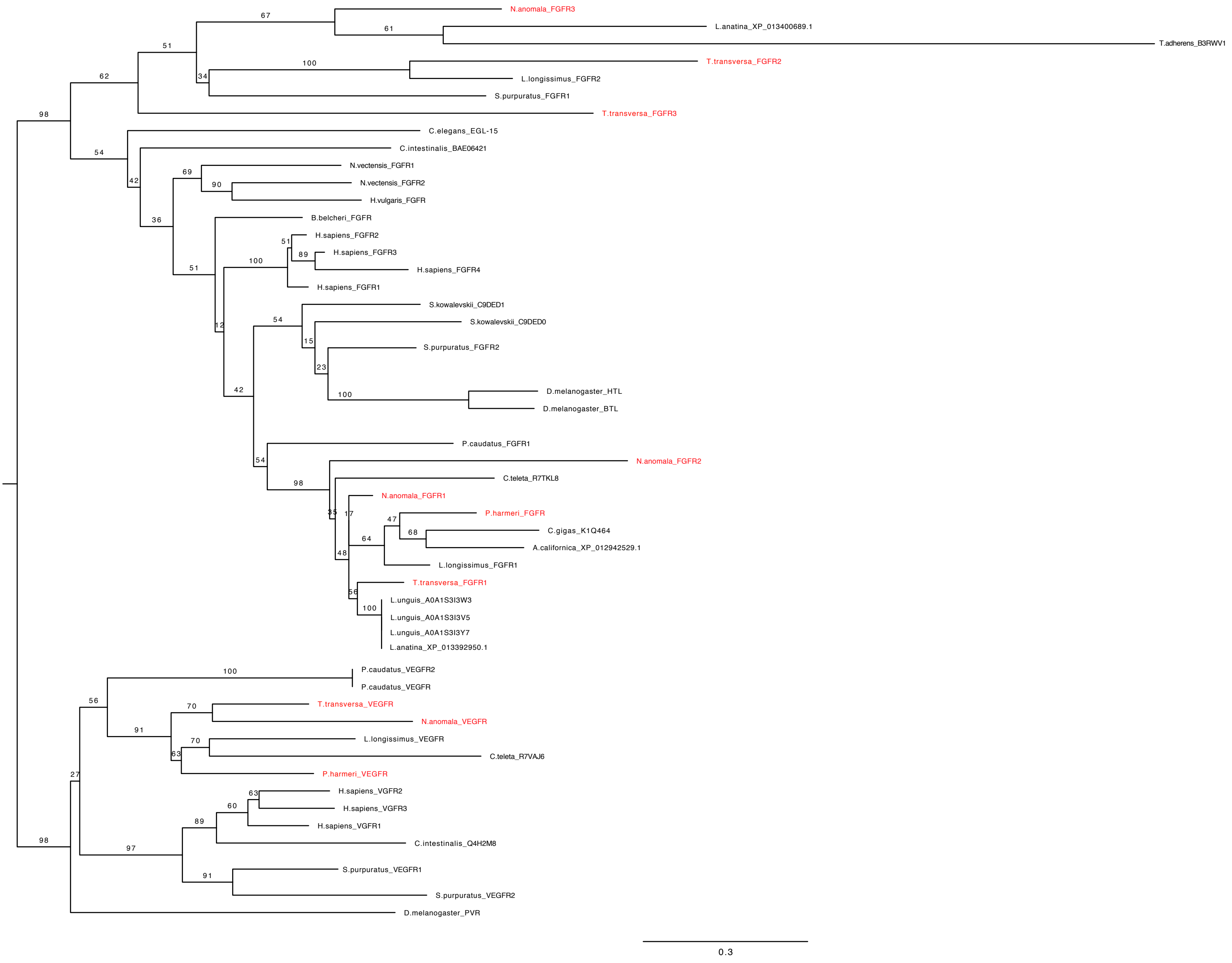

N.anomala_FGFR3
L.anatina_XP_013400689.1
T.adherens_B3RWV1
T.transversa_FGFR2
L.longissimus_FGFR2
S.purpuratus_FGFR1
T.transversa_FGFR3
C.elegans_EGL-15
C.intestinalis_BAE06421
N.vectensis_FGFR1
N.vectensis_FGFR2
H.vulgaris_FGFR
B.belcheri_FGFR
H.sapiens_FGFR2
H.sapiens_FGFR3
H.sapiens_FGFR4
H.sapiens_FGFR1
S.kowalevskii_C9DED1
S.kowalevskii_C9DED0
S.purpuratus_FGFR2
D.melanogaster_HTL
D.melanogaster_BTL
P.caudatus_FGFR1
N.anomala_FGFR2
C.teleta_R7TKL8
N.anomala_FGFR1
P.harmeri_FGFR
C.gigas_K1Q464
A.californica_XP_012942529.1
L.longissimus_FGFR1
T.transversa_FGFR1
L.unguis_A0A1S3I3W3
L.unguis_A0A1S3I3V5
L.unguis_A0A1S3I3Y7
L.anatina_XP_013392950.1
P.caudatus_VEGFR2
P.caudatus_VEGFR
T.transversa_VEGFR
N.anomala_VEGFR
L.longissimus_VEGFR
C.teleta_R7VAJ6
P.harmeri_VEGFR
H.sapiens_VGFR2
H.sapiens_VGFR3
H.sapiens_VGFR1
C.intestinalis_Q4H2M8
S.purpuratus_VEGFR1
S.purpuratus_VEGFR2
D.melanogaster_PVR
0.3


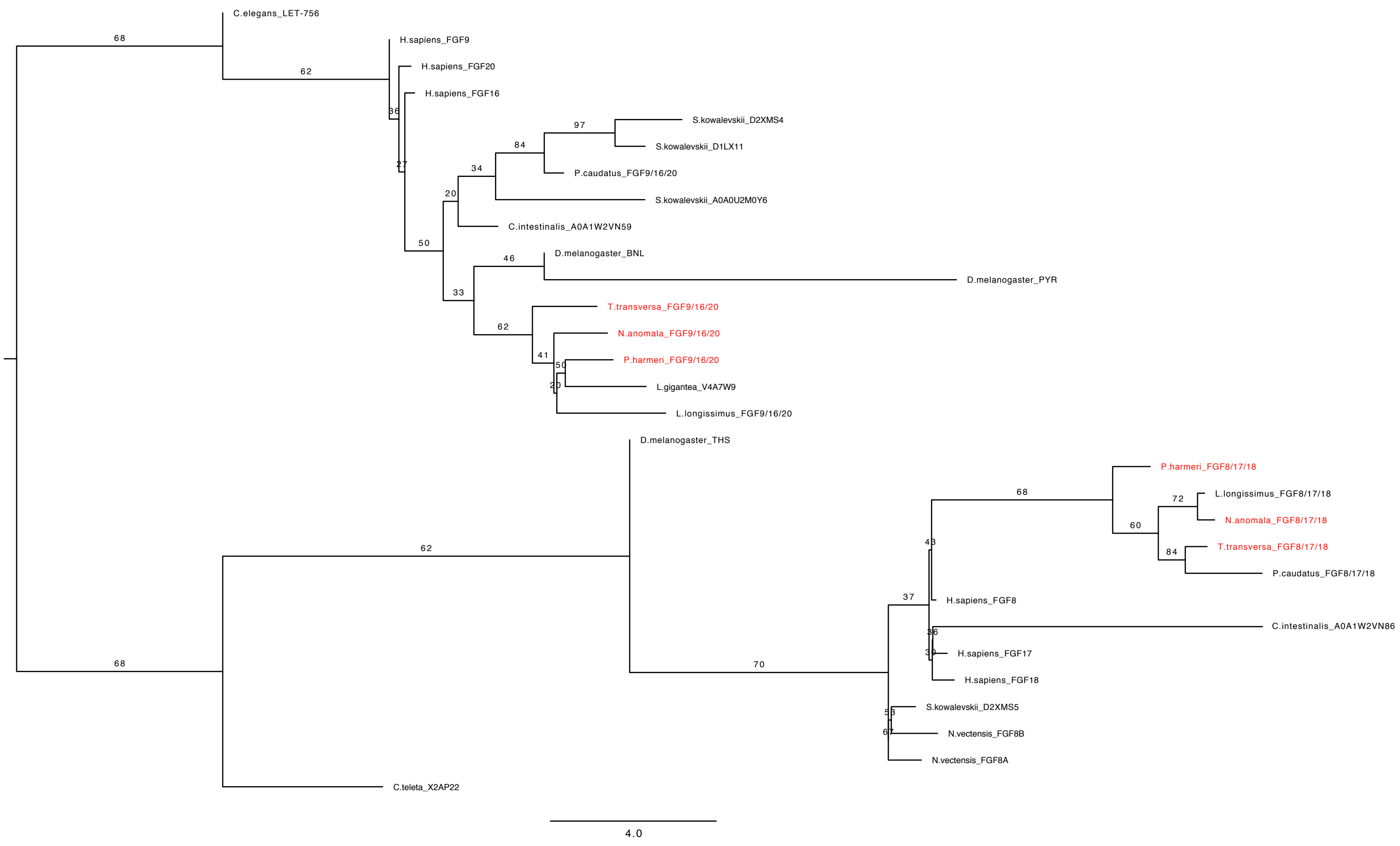

C.elegans_LET-756
H.sapiens_FGF9
H.sapiens_FGF20
H.sapiens_FGF16
S.kowalevskii_D2XMS4
S.kowalevskii_D1LX11
P.caudatus_FGF9/16/20
S.kowalevskii_A0A0U2M0Y6
C.intestinalis_A0A1W2VN59
D.melanogaster_BNL
D.melanogaster_PYR
T.transversa_FGF9/16/20
N.anomala_FGF9/16/20
P.harmeri_FGF9/16/20
L.gigantea_V4A7W9
L.longissimus_FGF9/16/20
D.melanogaster_THS
P.harmeri_FGF8/17/18
L.longissimus_FGF8/17/18
N.anomala_FGF8/17/18
T.transversa_FGF8/17/18
P.caudatus_FGF8/17/18
H.sapiens_FGF8
C.intestinalis_A0A1W2VN86
H.sapiens_FGF17
H.sapiens_FGF18
S.kowalevskii_D2XMS5
N.vectensis_FGF8B
N.vectensis_FGF8A
C.teleta_X2AP22
4.0

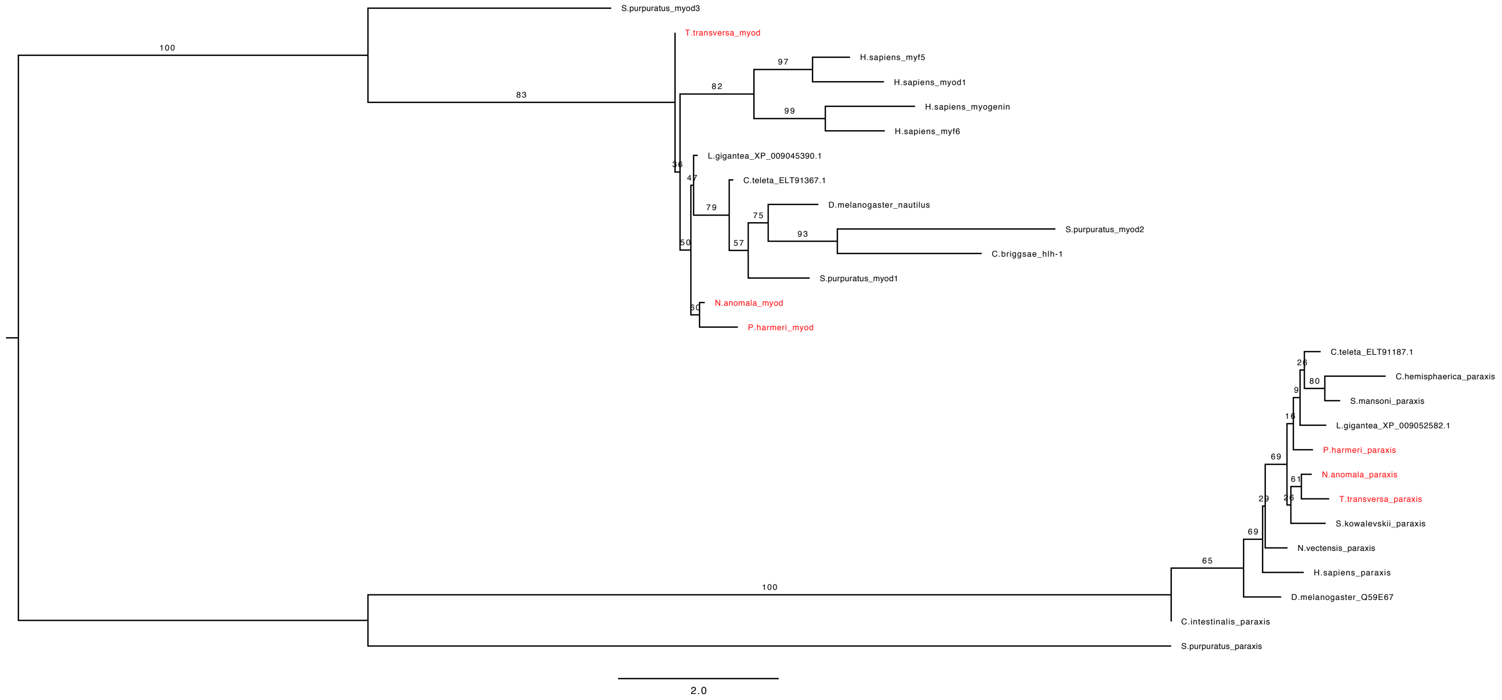
S.purpuratus_myod3
T.transversa_myod
H.sapiens_myf5
H.sapiens_myod1
H.sapiens_myogenin
H.sapiens_myf6
L.gigantea_XP_009045390.1
C.teleta_ELT91367.1
D.melanogaster_nautilus
S.purpuratus_myod2
C.briggsae_hlh-1
S.purpuratus_myod1
N.anomala_myod
P.harmeri_myod
C.teleta_ELT91187.1
C.hemisphaerica_paraxis
S.mansoni_paraxis
L.gigantea_XP_009052582.1
P.harmeri_paraxis
N.anomala_paraxis
T.transversa_paraxis
S.kowalevskii_paraxis
N.vectensis_paraxis
H.sapiens_paraxis
D.melanogaster_Q59E67
C.intestinalis_paraxis
S.purpuratus_paraxis
100
83
97
82
99
36
47
79
75
93
57
50
30
80
9
16
69
61
29
69
65
100
2.0
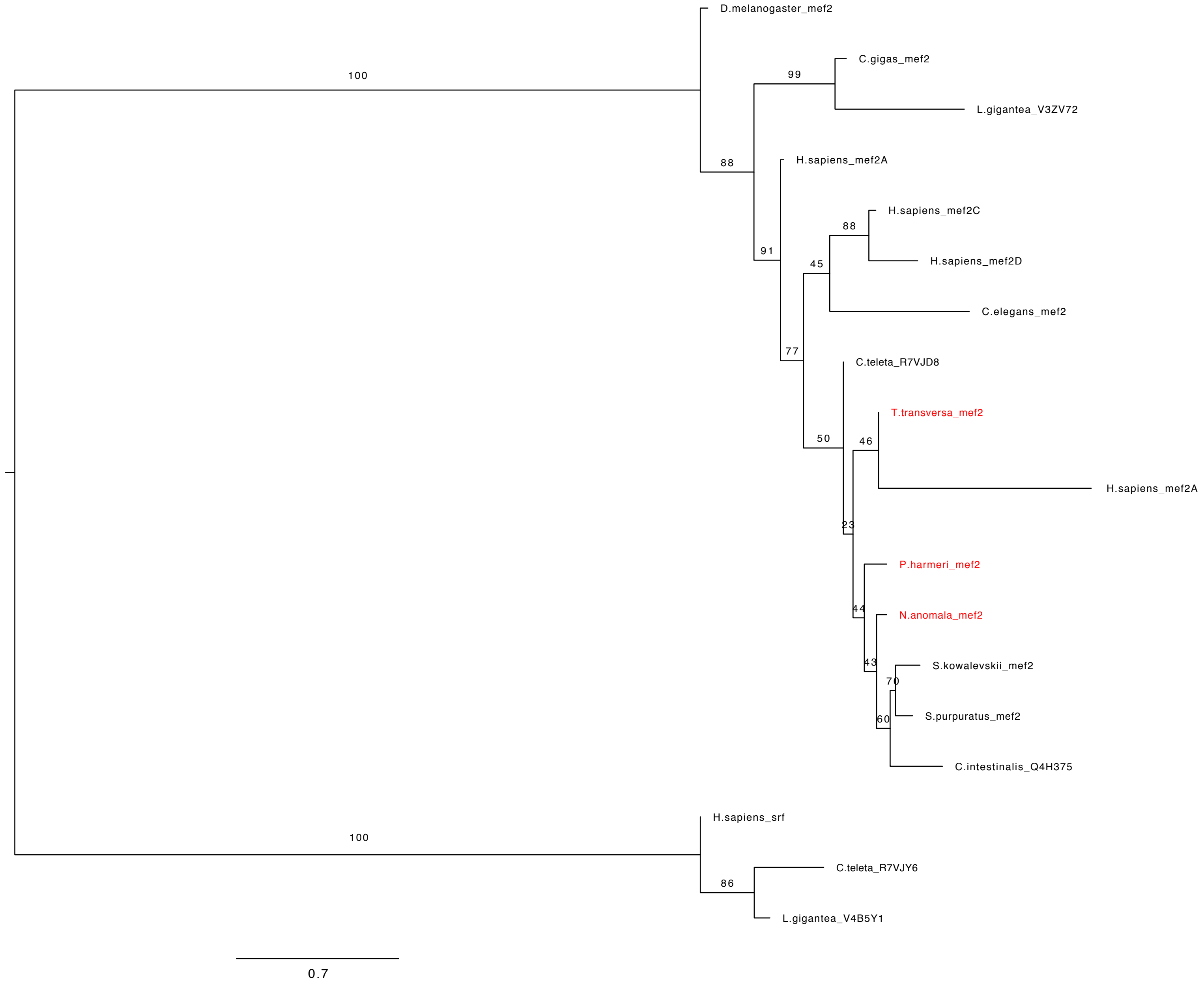
D.melanogaster_mef2
C.gigas_mef2
L.gigantea_V3ZV72
H.sapiens_mef2A
H.sapiens_mef2C
H.sapiens_mef2D
C.elegans_mef2
C.teleta_R7VJD8
T.transversa_mef2
H.sapiens_mef2A
P.harmeri_mef2
N.anomala_mef2
S.kowalevskii_mef2
S.purpuratus_mef2
C.intestinalis_Q4H375
H.sapiens_srf
C.teleta_R7VJY6
L.gigantea_V4B5Y1
100
99
88
91
88
45
77
50
46
23
44
43
70
60
100
86
0.7

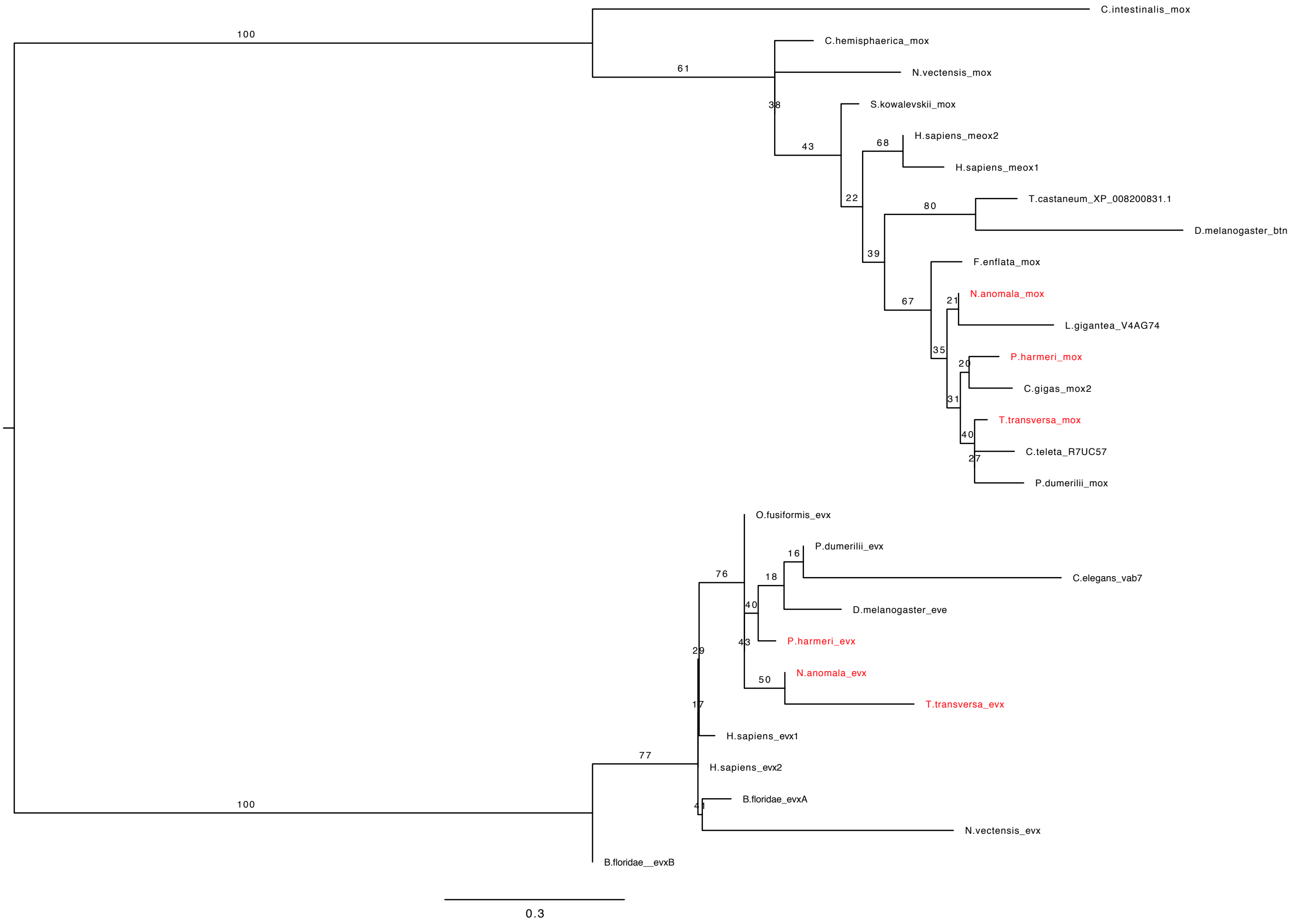
C.intestinalis_mox
C.hemisphaerica_mox
N.vectensis_mox
S.kowalevskii_mox
H.sapiens_meox2
H.sapiens_meox1
T.castaneum_XP_008200831.1
D.melanogaster_btn
F.enflata_mox
N.anomala_mox
L.gigantea_V4AG74
P.harmeri_mox
C.gigas_mox2
T.transversa_mox
C.teleta_R7UC57
P.dumerilii_mox
O.fusiformis_evx
P.dumerilii_evx
C.elegans_vab7
D.melanogaster_eve
P.harmeri_evx
N.anomala_evx
T.transversa_evx
H.sapiens_evx1
H.sapiens_evx2
B.floridae_evxA
N.vectensis_evx
B.floridae__evxB
0.3

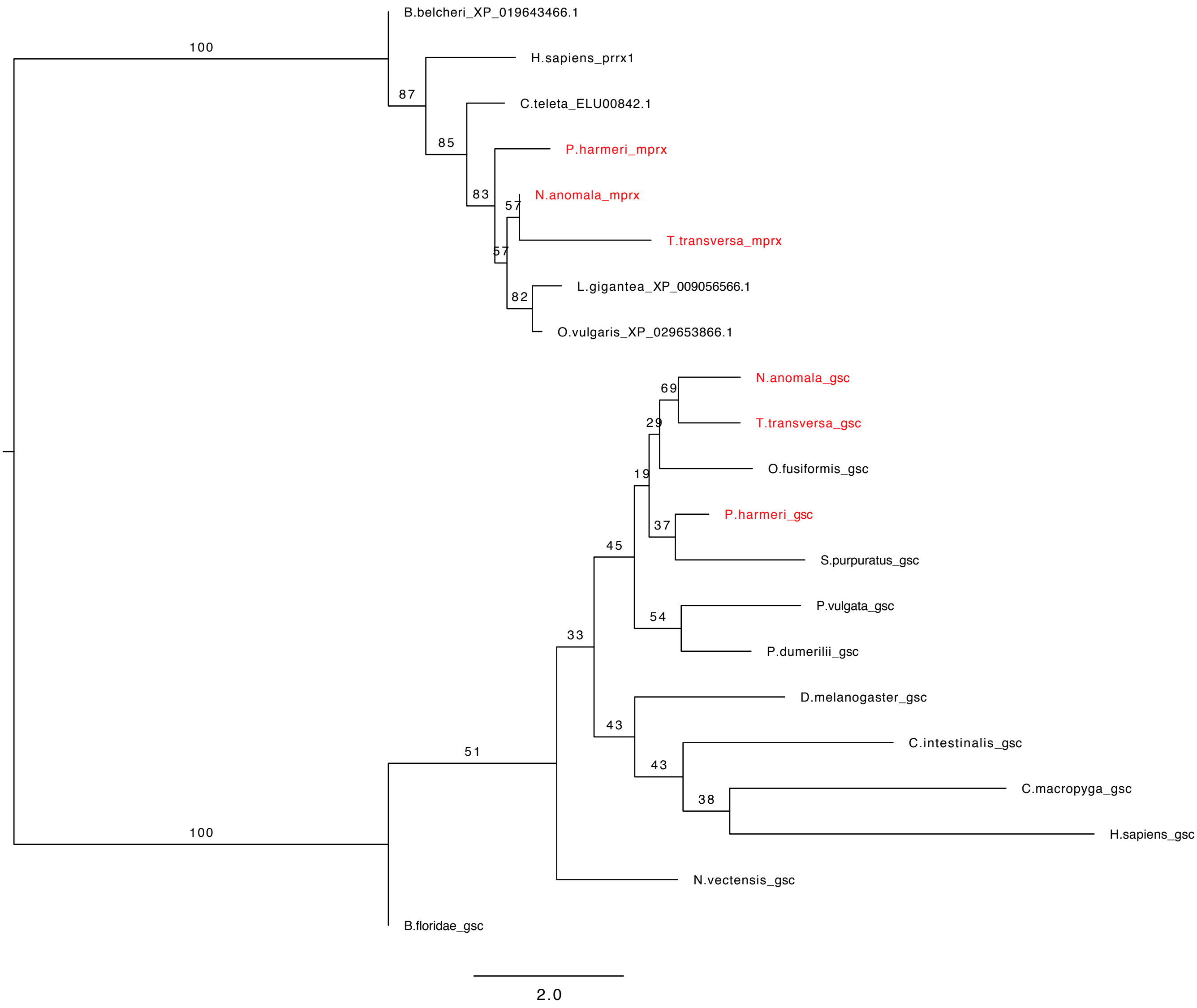
B.belcheri_XP_019643466.1
H.sapiens_prrx1
C.teleta_ELU00842.1
P.harmeri_mprx
N.anomala_mprx
T.transversa_mprx
L.gigantea_XP_009056566.1
O.vulgaris_XP_029653866.1
N.anomala_gsc
T.transversa_gsc
O.fusiformis_gsc
P.harmeri_gsc
S.purpuratus_gsc
P.vulgata_gsc
P.dumerilii_gsc
D.melanogaster_gsc
C.intestinalis_gsc
C.macropyga_gsc
H.sapiens_gsc
N.vectensis_gsc
B.floridae_gsc
2.0

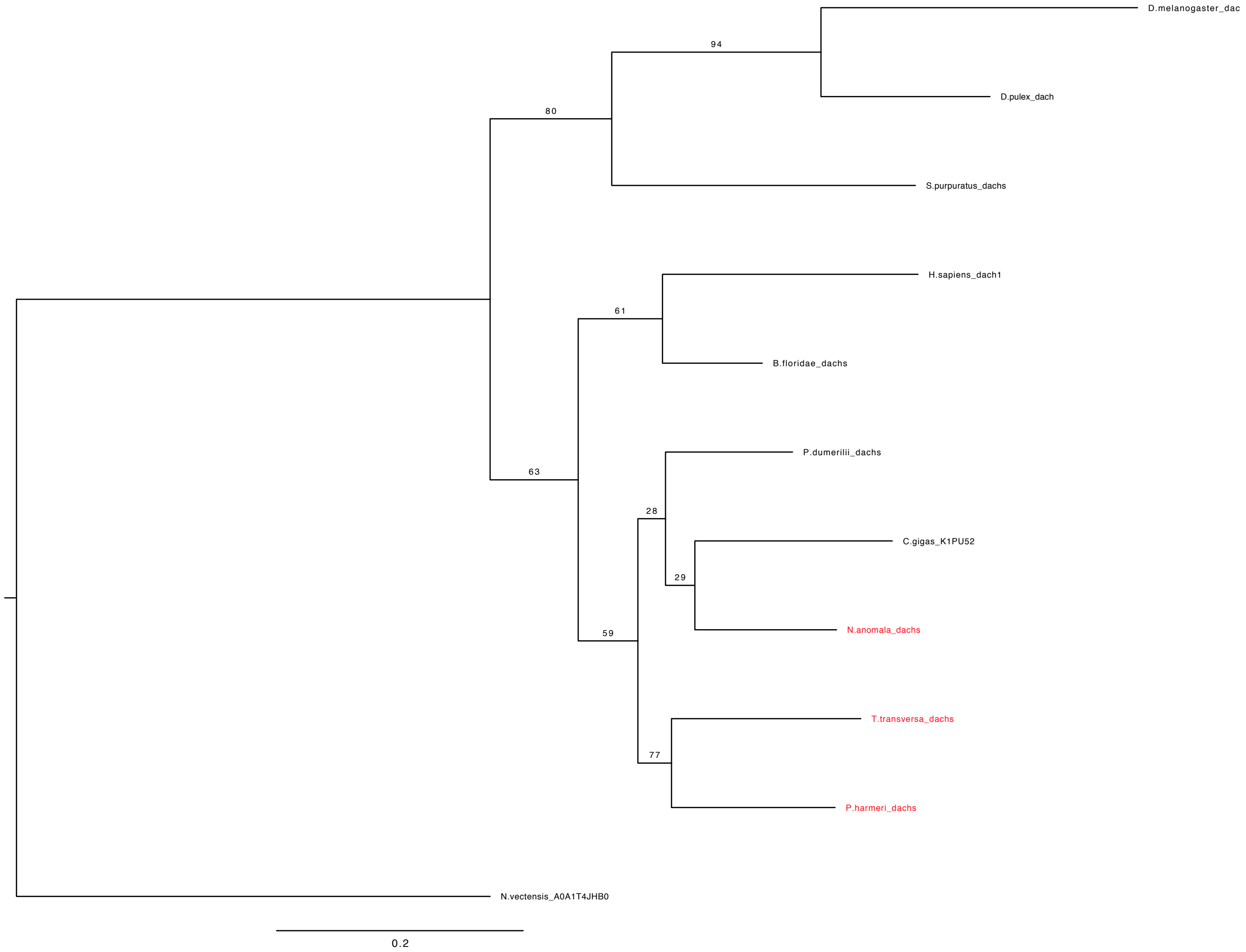
D.melanogaster_dac
94
D.pulex_dach
80
S.purpuratus_dachs
H.sapiens_dach1
61
B.floridae_dachs
P.dumerilii_dachs
63
28
C.gigas_K1PU52
29
N.anomala_dachs
59
T.transversa_dachs
77
P.harmeri_dachs
N.vectensis_A0A1T4JHB0
0.2

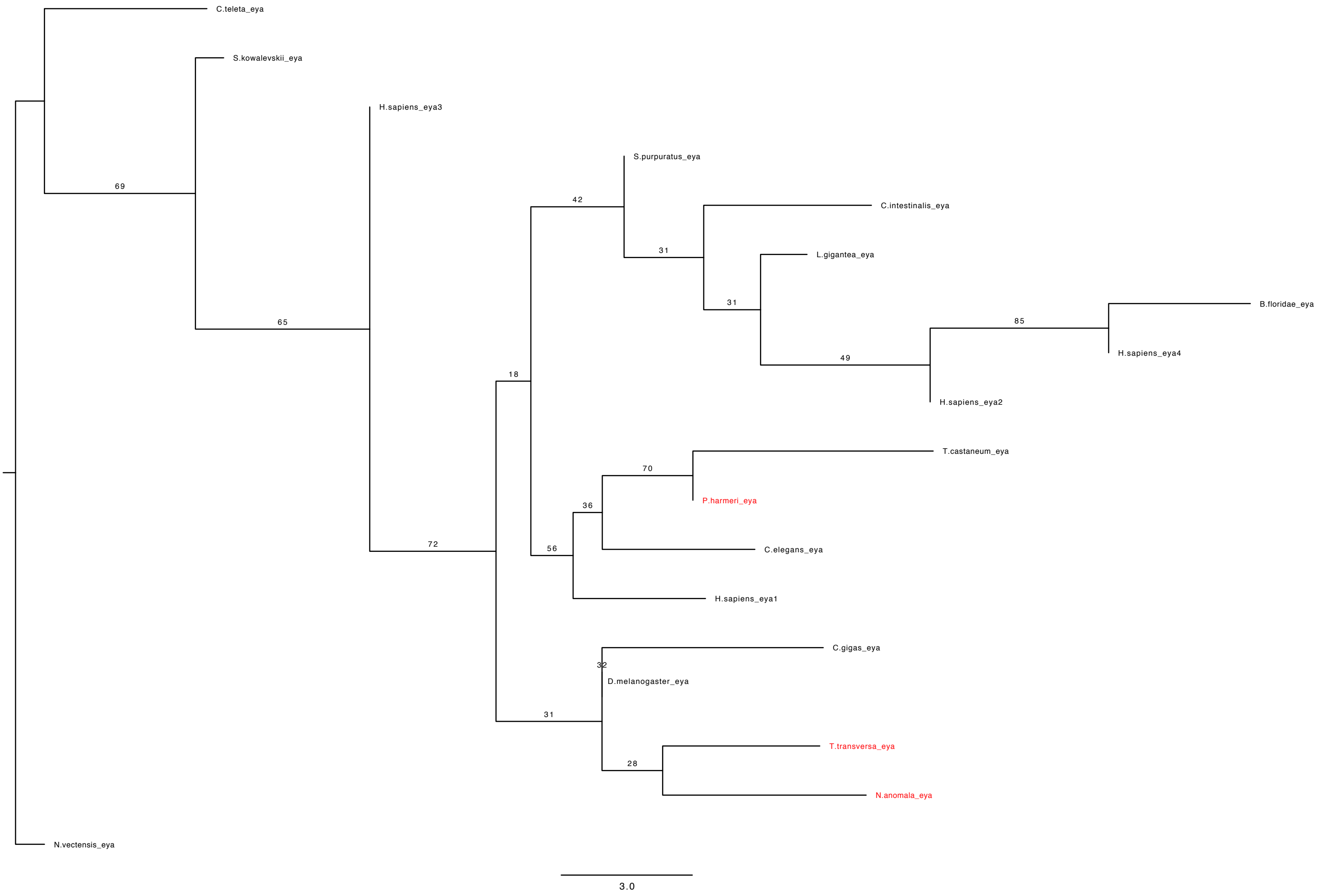
C.teleta_eya
S.kowalevskii_eya
H.sapiens_eya3
69
S.purpuratus_eya
42
C.intestinalis_eya
31
L.gigantea_eya
31
B.floridae_eya
85
65
H.sapiens_eya4
49
H.sapiens_eya2
18
T.castaneum_eya
70
P.harmeri_eya
36
72
56
C.elegans_eya
H.sapiens_eya1
C.gigas_eya
32
D.melanogaster_eya
31
T.transversa_eya
28
N.anomala_eya
N.vectensis_eya
3.0

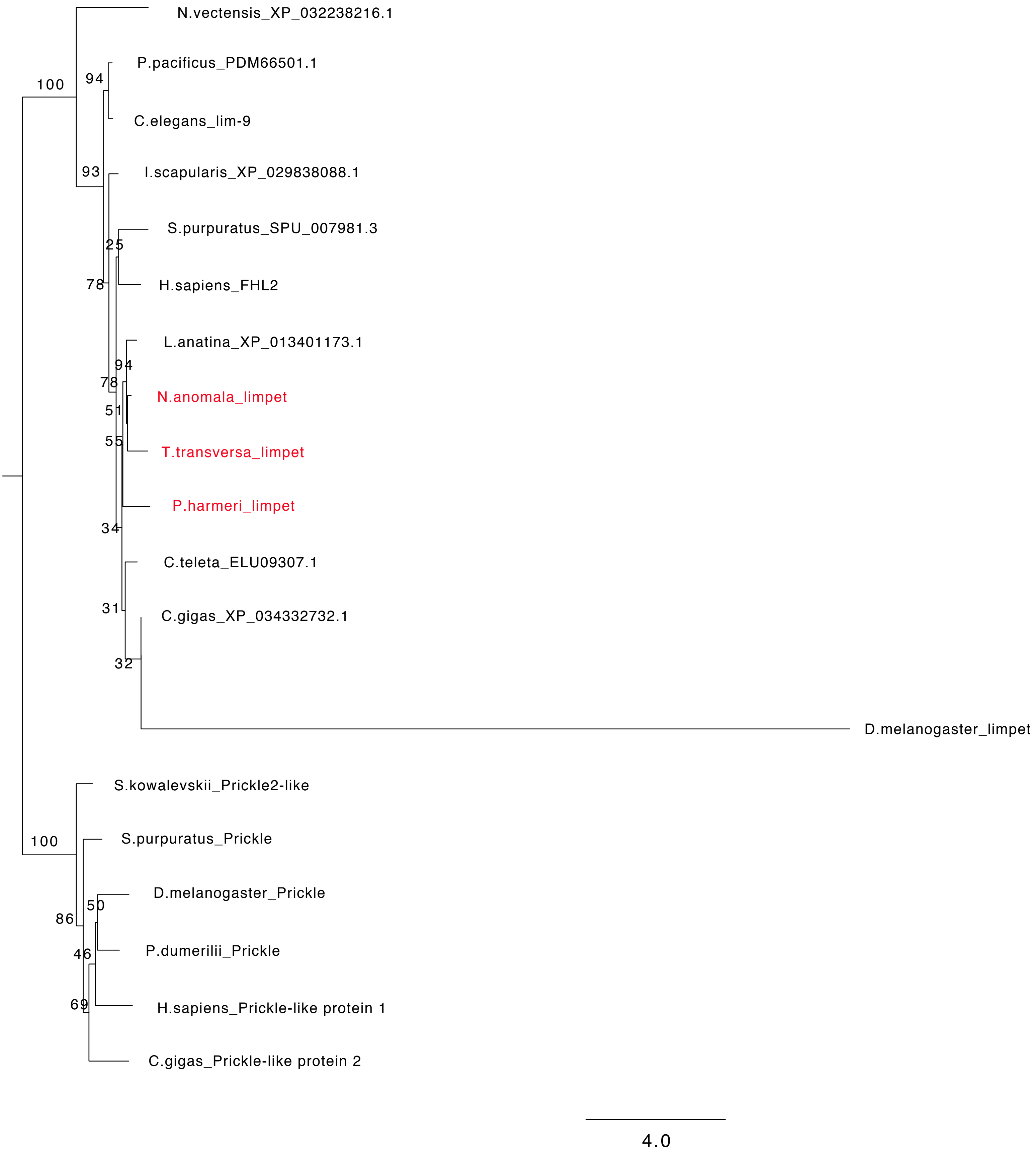
N.vectensis_XP_032238216.1
P.pacificus_PDM66501.1
C.elegans_lim-9
I.scapularis_XP_029838088.1
S.purpuratus_SPU_007981.3
H.sapiens_FHL2
L.anatina_XP_013401173.1
N.anomala_limpet
T.transversa_limpet
P.harmeri_limpet
C.teleta_ELU09307.1
C.gigas_XP_034332732.1
D.melanogaster_limpet
S.kowalevskii_Prickle2-like
S.purpuratus_Prickle
D.melanogaster_Prickle
P.dumerilii_Prickle
H.sapiens_Prickle-like protein 1
C.gigas_Prickle-like protein 2
100
94
93
25
78
94
78
51
55
34
31
32
100
50
86
46
69
4.0

Fig. S1. **Orthology analysis.** Putative orthologous sequences were identified by tBLASTx search of the transcriptome of *T. transversa, N. anomala* and *Ph. harmeri*. Bayesian phylogenetic analysis is supporting orthology. Names of genes or proteins, if available, follow the name of organism(s). *T. transversa, N. anomala* and *Ph. harmeri* sequences are highlighted in red.

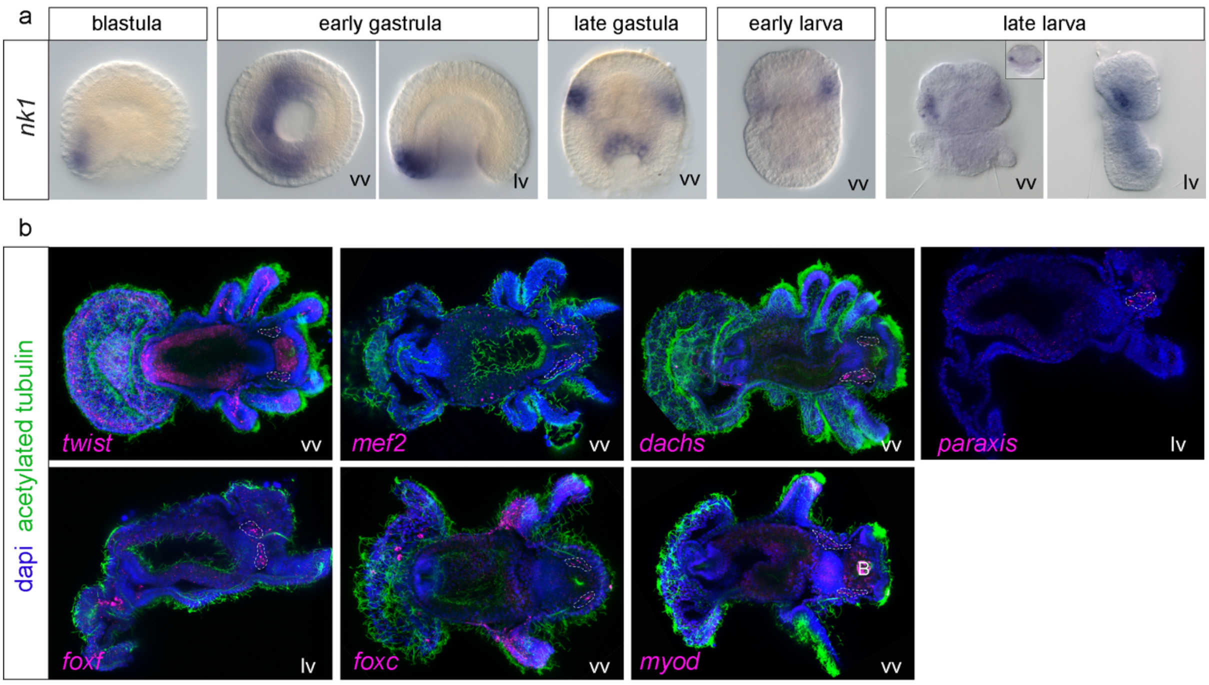


Fig. S2. **Additional gene expression patterns**. a) WMISH of *nk1* in blastulae, early gastrulae, late gastrulae, early larvae and late larvae of *N. anomala*. Anterior to the top. b) Fluorescent WMISH of *twist, mef2, dachs, paraxis, foxf, foxc* and *myod* in the 6-tentacle larva stage of *Ph. harmeri*. Gene expression is in magenta, cilia are labeled green with anti-acetylated tubulin antibody and nuclei are stained blue with DAPI. Posterior mesoderm is encircled with a white dashed line. Anterior to the left. lv, lateral view; vv, vegetal view. B, background staining.

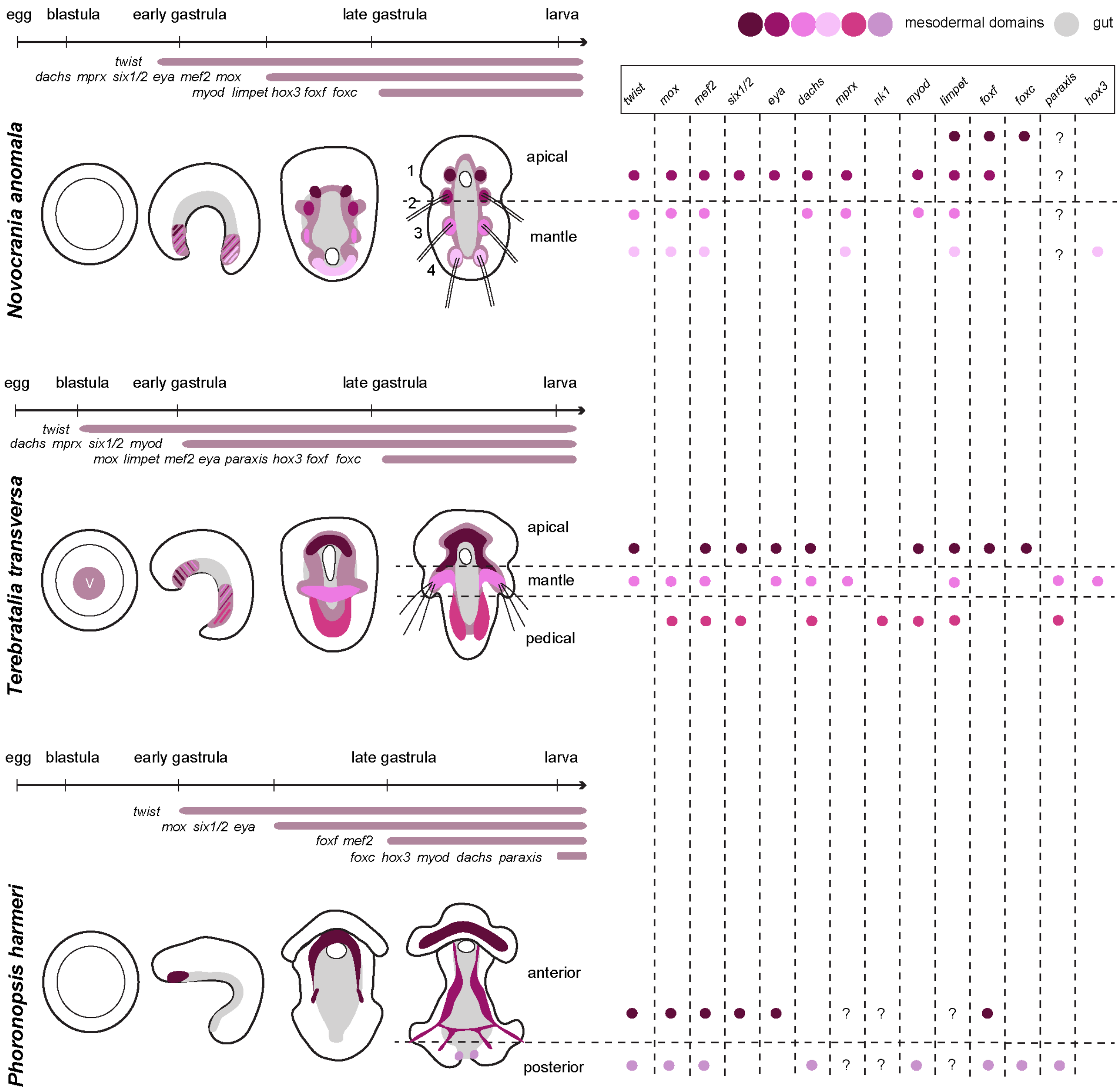


Fig. S3. **Comparison of mesodermal gene expression patterns in representative developmental stages of *N. anomala, T. transversa* and *Ph. harmeri*.** Schematic representation of the temporal and spatial expression patterns of *twist, mox, mef2, six1/2, eya, dachs, mprx, nk1, myod, limpet, foxf, foxc* and *hox3* in blastula, mid gastrula and larva stages of *N. anomala, T. transversa* and *Ph. harmeri*. *Nk1* and *hox3* do not have a mesodermal expressed in *N. anomala* and *Ph. harmeri,* respectively. Question marks indicates that the expression of these genes is not yet known. Drawings are not up to scale. Anterior to the top. V, vegetal view; A, animal view.

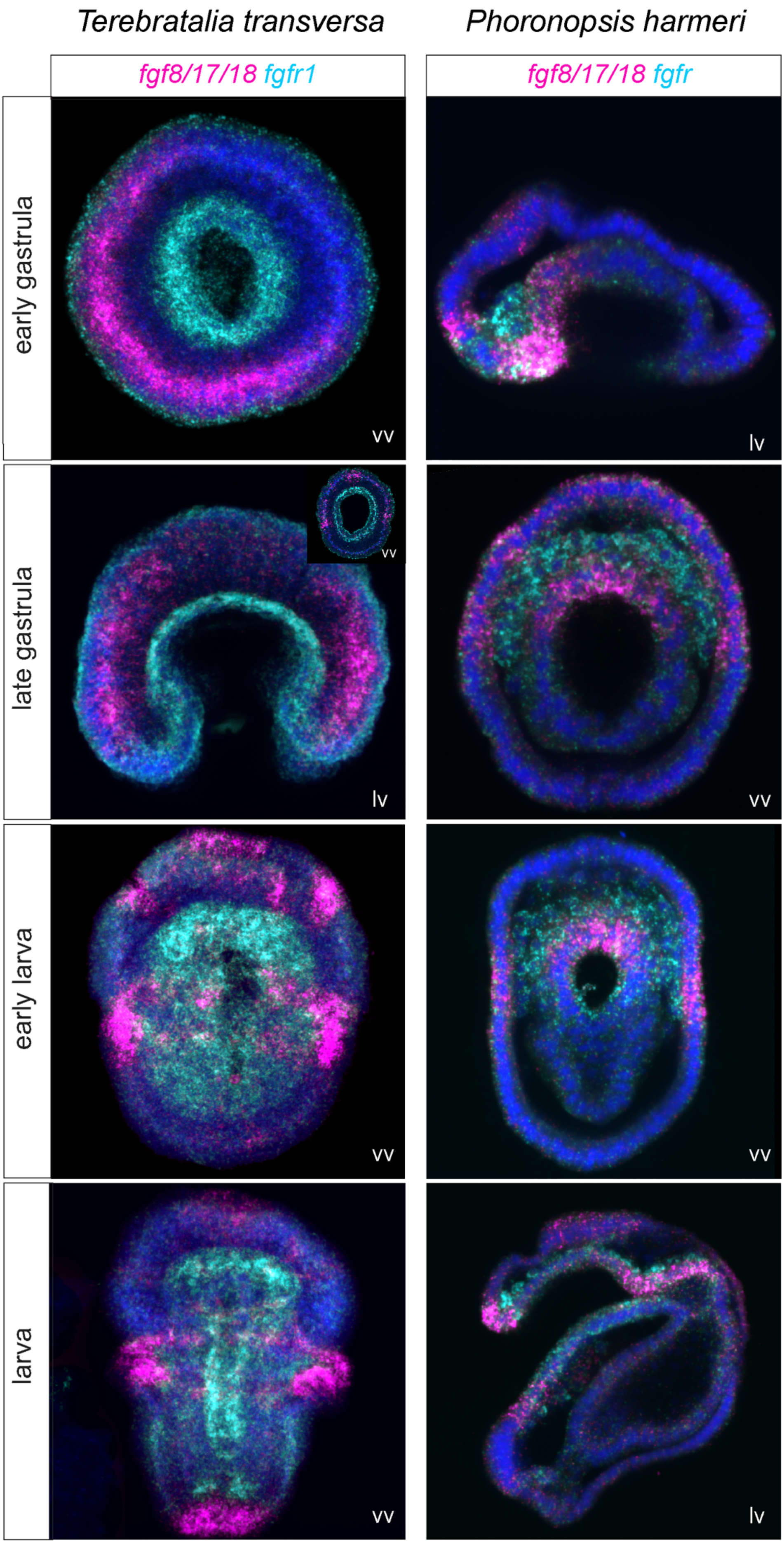

Terebratalia transversa
Phoronopsis harmeri
fgf8/17/18 fgfr1
fgf8/17/18 fgfr
early gastrula
late gastrula
early larva
larva
vv
lv

Fig. S4. **Co-expression analysis of *fgfr* (cyan) and *fgf8/11/18* (magenta) by double fluorescent WMISH in *T. transversa* and *Ph. harmeri*.** The inset shows an embryo in vegetal view. Every fluorescent image is a full projection of merged confocal stacks and nuclei are stained with DAPI. Anterior to the top. lv, lateral view; vv, vegetal view.

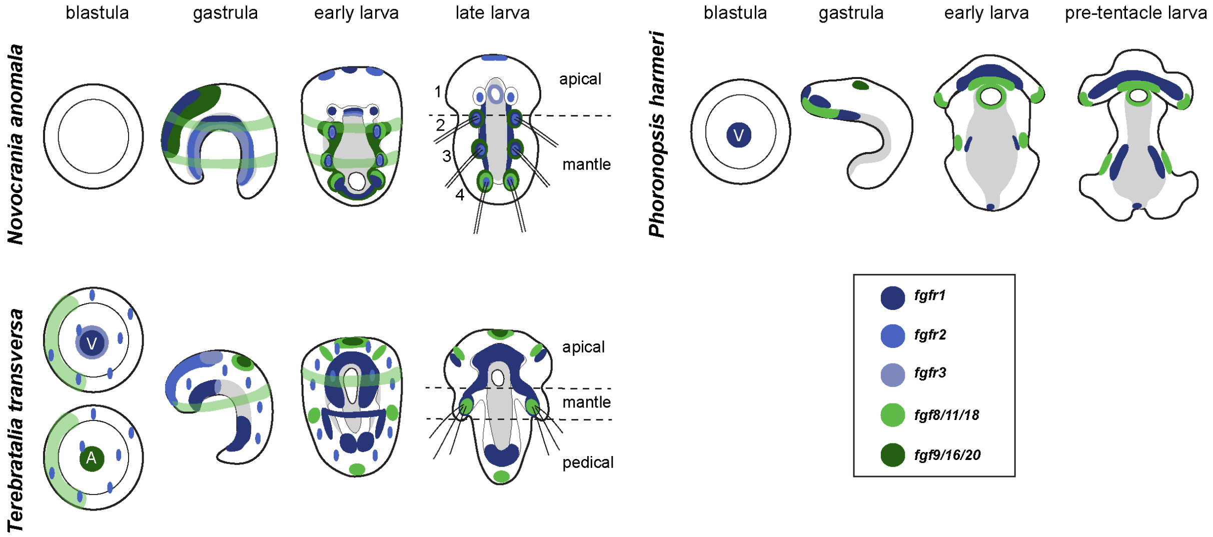


Fig. S5. **Comparison of gene expression patterns of the FGF signaling components in representative developmental stages of *Novocrania anomala, Terebratalia transversa* and *Ph. harmeri*.** Schematic representation of the expression patterns of *fgfr1, fgfr2, fgfr3, fgf8/11/18* and *fgf9/16/20* in blastula, mid gastrula and larva stages of *N. anomala, T. transversa* and *Ph. harmeri*. Drawings are not up to scale. Anterior to the top. V, vegetal view; A, animal view.

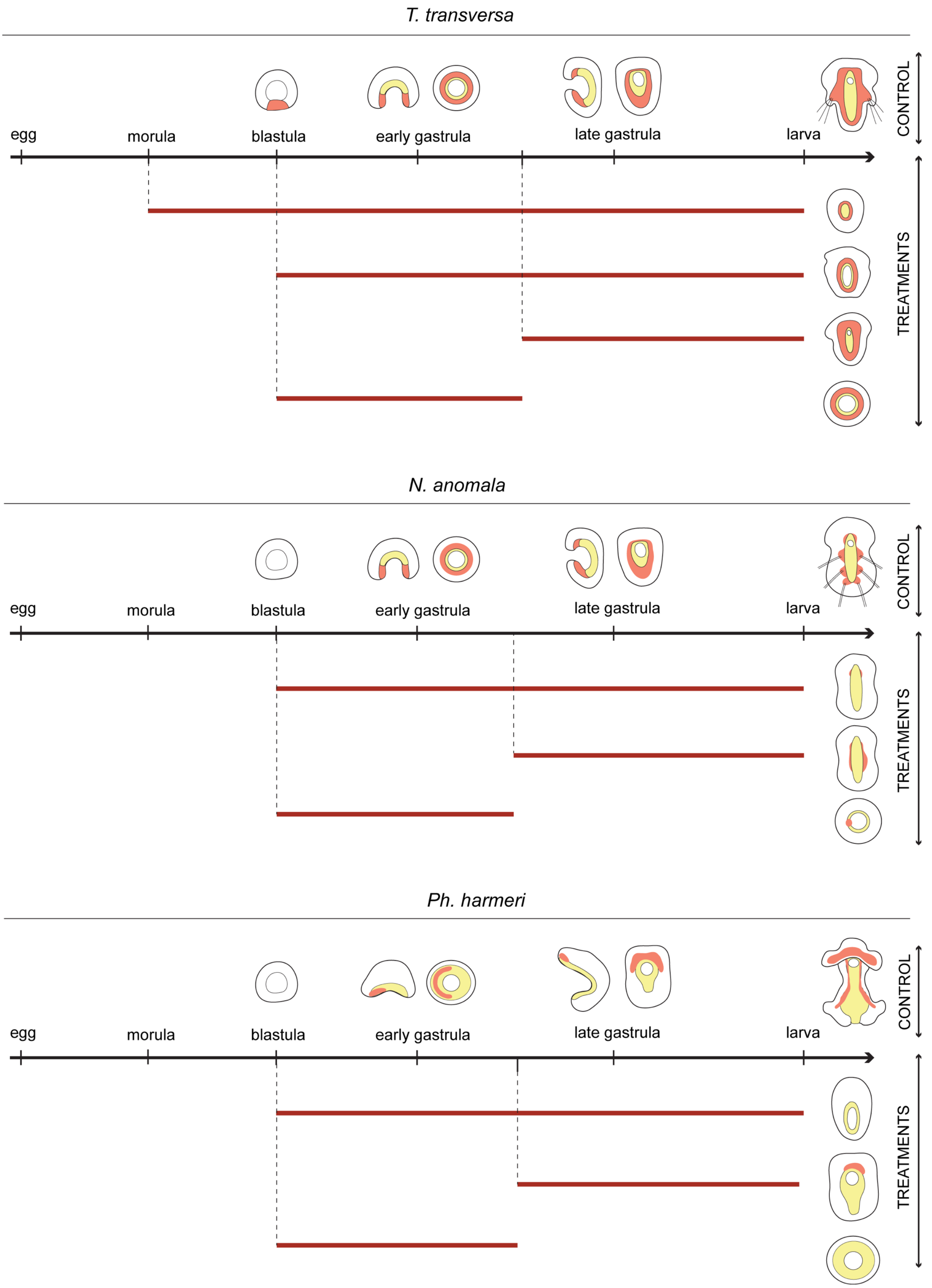


Fig. S6. **Summary of 20 uM SU5402 treatments and observed phenotypes in *T. transversa, N. anomala* and *Ph. harmeri*.** Embryos were treated with 20 uM SU5402 from morula (only in *T. transversa*), blastula and gastrula stages and fixed at larva stage. Embryos treated from blastula stage were also fixed at gastrula stage. Anterior to the top. Drawings are not up to scale.

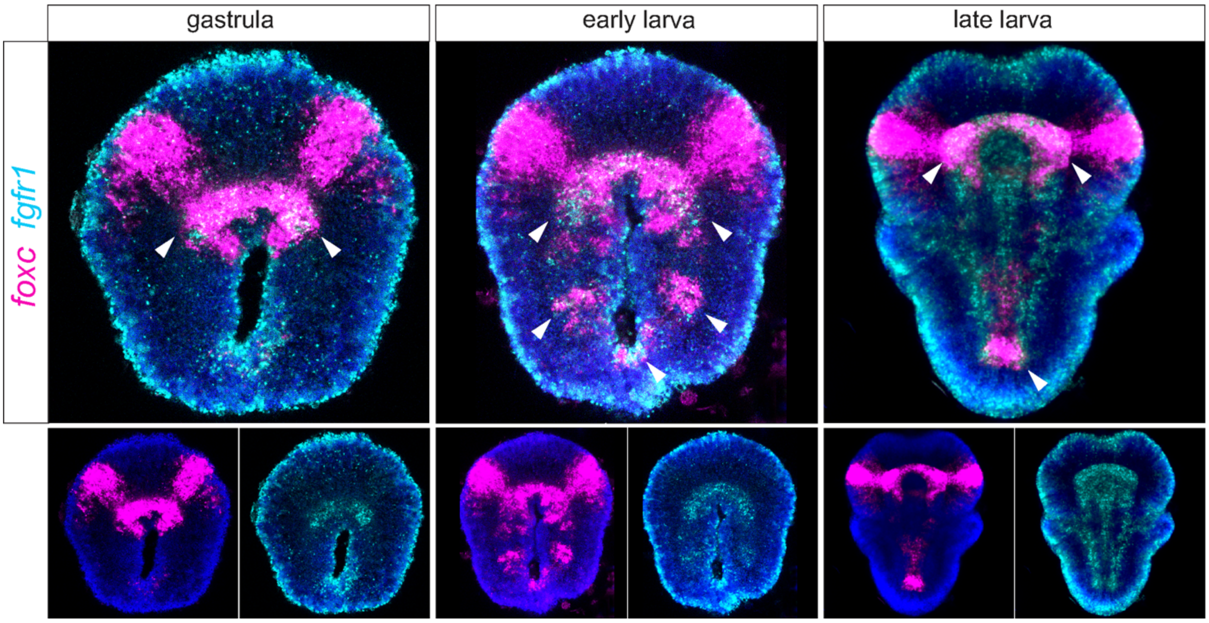


Fig. S7. **Co-expression analysis of *fgfr* (cyan) and *foxc* (magenta) by double fluorescent WMISH in gastrula and larva stages of *T. transversa*.** White arrowheads indicate co-expression. Every fluorescent image is a full projection of merged confocal stacks and nuclei are stained with DAPI. Anterior to the top. All panels depict embryos in vegetal view.

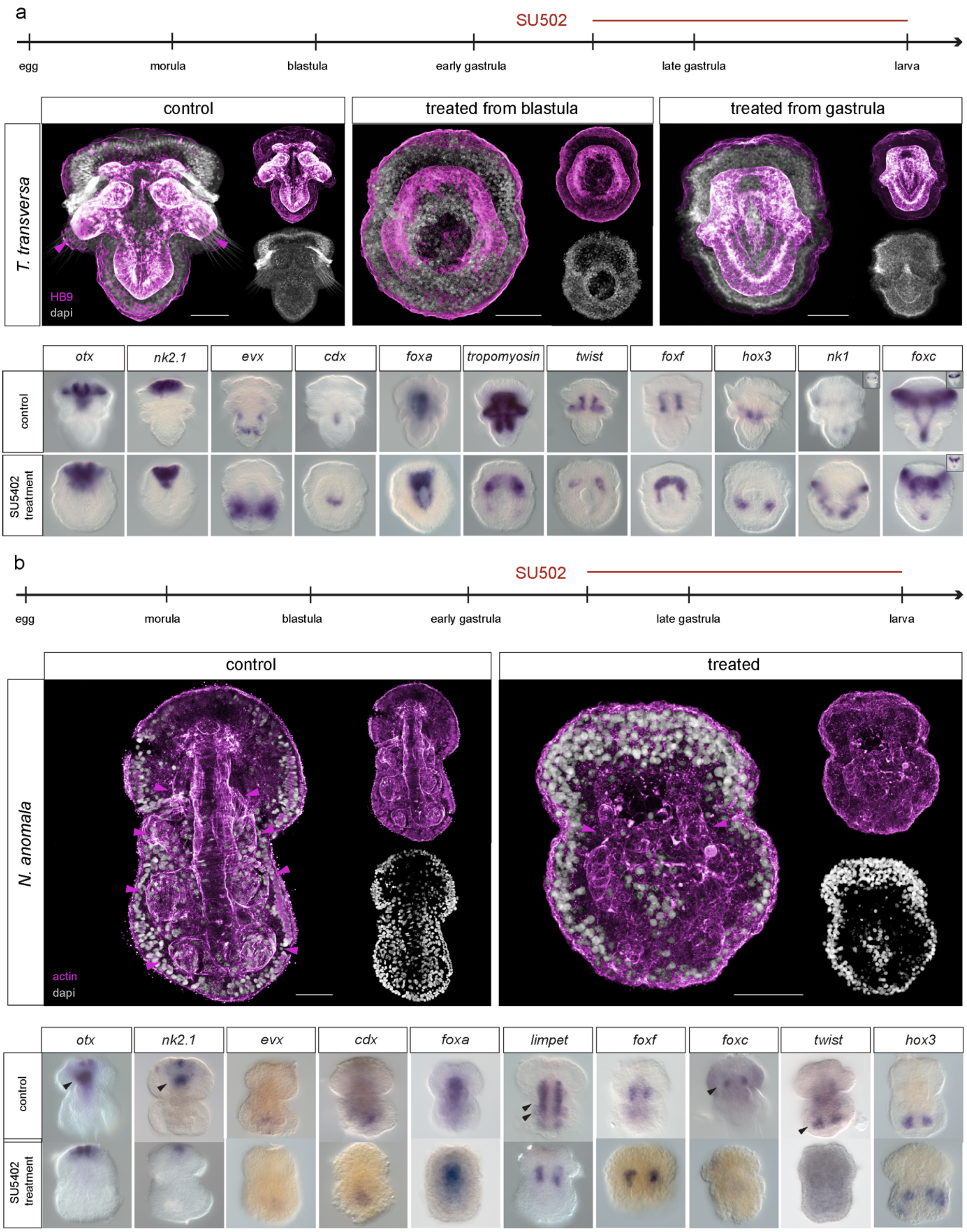


Fig. S8. **Additional SU5402 treatments in brachiopods.** (a) Immunohistochemistry of HB9, which labels mesoderm in *T. transversa* blastula and gastrula embryos treated with 20 uM SU5402 and fixed at larva stage. Magenta arrowheads indicate mesodermal domains associated with chaetae sacs. Every fluorescent image is a full projection of merged confocal stacks and nuclei are stained with DAPI. WMISH of anterior (*otx, nk2.1*), posterior *(evx)* and endodermal *(foxa, cdx)* genes, musculature (*tropomyosin)*, anterior mesoderm (*twist*, *foxf),* mid mesoderm (*hox3*) and posterior mesoderm *(nk1, foxc)* in *T. transversa* gastrula embryos treated with 20 uM SU5402 and fixed at larva stage. Insets show different focal planes of the embryos. (b) Immunohistochemistry of musculature (actin) in *N. anomala* gastrula embryos treated with 20 uM SU5402 and fixed at larva stage. Magenta arrowheads show the musculature associated to coelomic and chaetae sacs. WMISH of anterior (*otx, nk2.1*), posterior genes (*evx*) and endodermal *(foxa, cdx)* genes*,* the whole mesoderm (*limpet)*, anterior mesoderm (*foxf, foxc)* and posterior mesoderm *(twist, hox3)*, in *N. anomala* gastrula embryos treated with 20 uM SU5402 and fixed at larva stage. Black arrowheads indicate the domains of expression that are absent in the treated embryos. All panels depict embryos in vegetal view. Anterior to the top. Scale bar: 20 um.

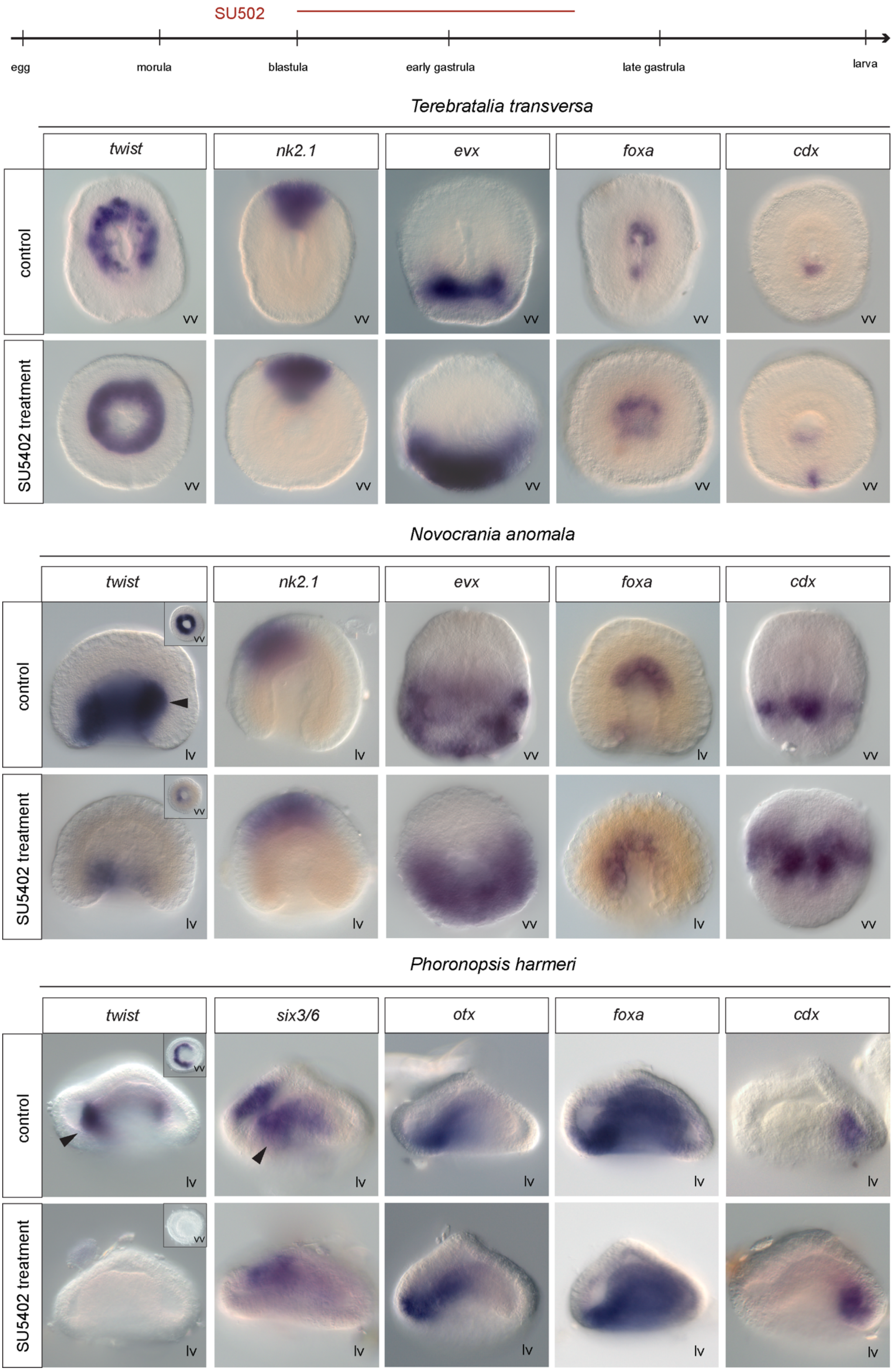

SU502
egg
morula
blastula
early gastrula
late gastrula
larva
Terebratalia transversa
twist
nk2.1
evx
foxa
cdx
control
SU5402 treatment
vv
Novocrania anomala
twist
nk2.1
evx
foxa
cdx
control
SU5402 treatment
lv
vv
Phoronopsis harmeri
twist
six3/6
otx
foxa
cdx
control
SU5402 treatment
lv
vv

Fig. S9. **WMISH on SU5402 treated *T. transversa, N. anomala* and *Ph. harmeri* embryos fixed at gastrula stage.** WMISH of mesodermal (*twist, six3/6),* endodermal *(foxa),* anterior (*otx*, *nk2.1, six3/6*) and posterior genes (*evx, cdx*) in *T. transversa, N. anomala* and *Ph. harmeri* blastula embryos treated with 20 uM SU5402 and fixed at gastrula stage. Black arrowheads indicate the domains of expression that are absent in the treated embryos. Insets show different focal planes of the embryos. Anterior to the top. lv, lateral view; vv, vegetal view.

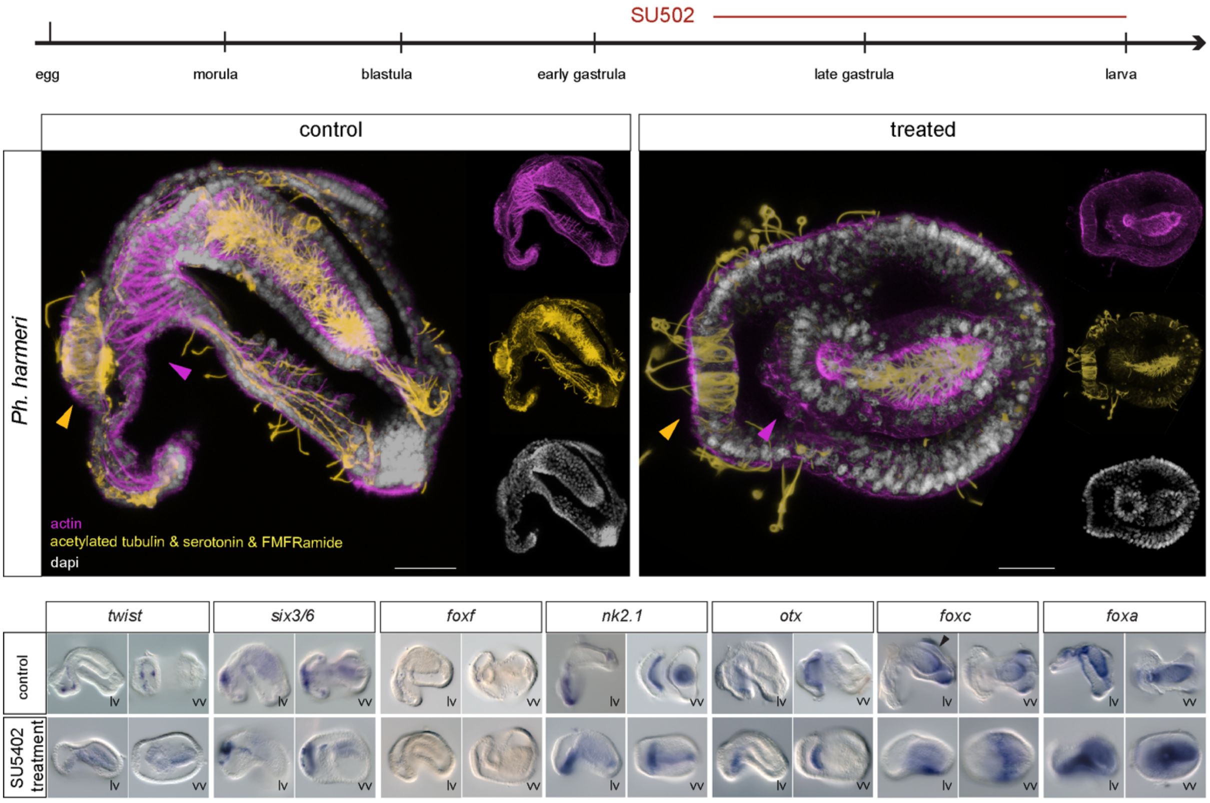


Fig. S10. **Additional SU5402 treatments in *Ph. harmeri.*** Immunohistochemistry of markers of the nervous system (acetylated tubulin, serotonin, FMFRamide) and musculature (actin) in *Ph. harmeri* gastrula embryos treated with 20 uM SU5402 and fixed at larva stage. WMISH of mesodermal (*twist*, *six3/6, foxf*), anterior (*six3/6, otx, nk2.1*), postero-ventral (*foxc*) and endodermal (*foxa, nk2.1*) genes in *Ph. harmeri* gastrula embryos treated with 20 uM SU5402 and fixed at larva stage. Yellow arrowheads indicate the apical organ and magenta arrowheads the esophageal musculature. Black arrowheads indicate the domains of expression that are absent in the treated embryos. Every fluorescent image is a full projection of merged confocal stacks and nuclei are stained with DAPI. Anterior to the left. lv, lateral view; vv, vegetal view. Scale bar: 20 um.

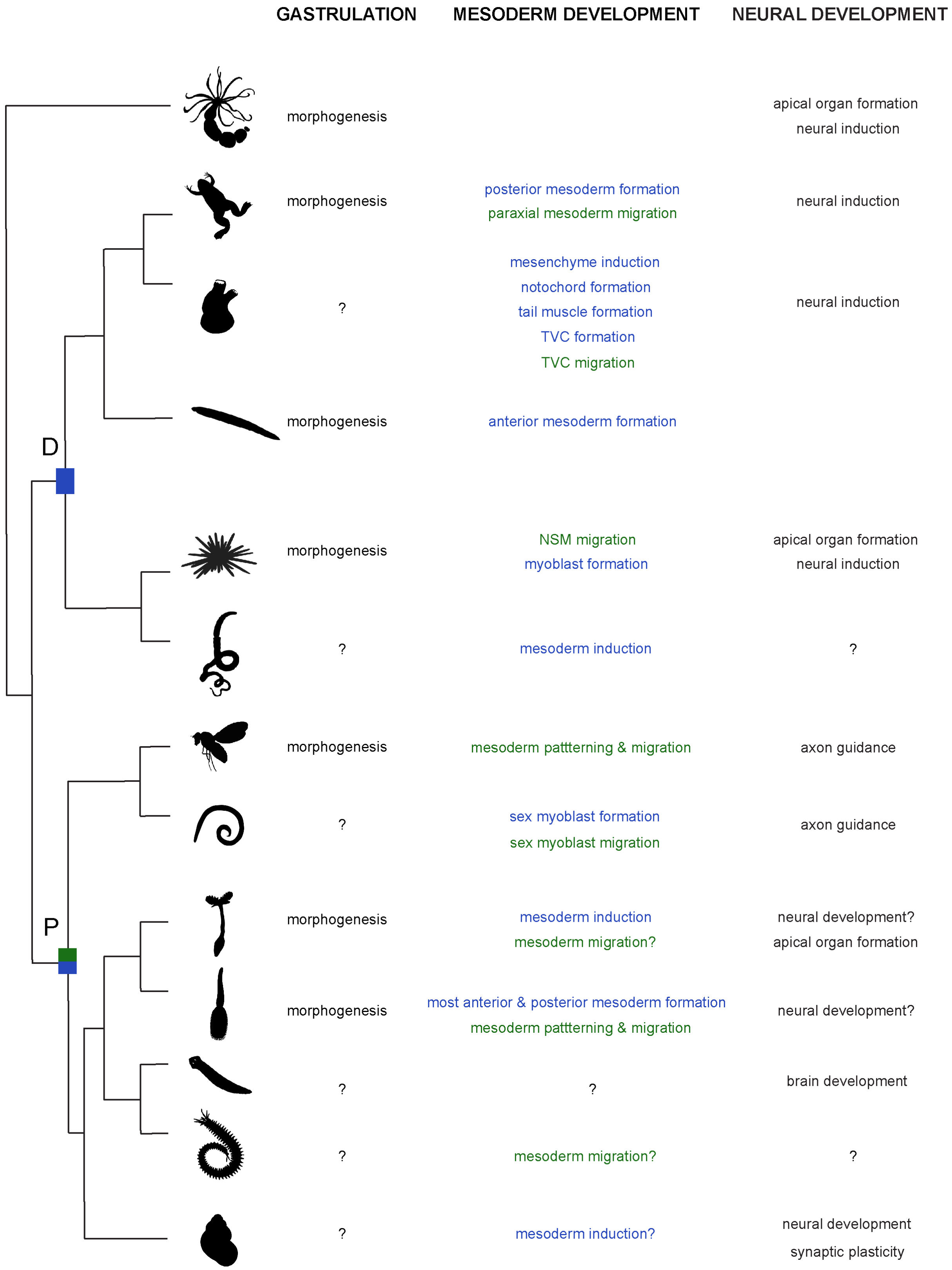


Fig. S11. **The role of FGF signaling in investigated species.** Table summarizing case studies where FGF signaling is found to be upstream of mesoderm development, morphogenetic movements of gastrulation and neural development. Animal illustrations are taken from phylopic.org (CC BY 3.0). D, deuterostomes; NSM, non-skeletogenic mesoderm; P, protostomes; TVC, trunk ventral cells.
